# SLAMF1-peptide mediated epigenetic priming reprograms innate immune responses in sepsis

**DOI:** 10.64898/2025.12.29.696918

**Authors:** Sindre Ullmann, Birgitta Ehrnström, Jørgen Stenvik, Sneha Pinto, Yashwanth Subbannayya, Victor Boyartchuk, Ingvild Bergdal Mestvedt, Mahamaya Dhaware, Siddhesh S. Kamat, Liv Ryan, Hilde Vagle, Tuva Børresdatter Dahl, Bente Evy Halvorsen, Jan Kristian Damås, Terje Espevik, Maria Yurchenko

**Author notes:** Corresponding author: **Address**: NTNU, Department of Clinical and Molecular Medicine (IKOM), P.O. Box 8905, 7491 Trondheim, Norway. **E-mail:**.

## Abstract

Sepsis is characterized by profound immune dysregulation, including impaired innate immune responses and epigenetic reprogramming of monocytes. However, strategies to therapeutically restore immune function remain limited. Here, we identify a cell-penetrating peptide, P7-Pen, as a modulator of monocyte epigenetic state and inflammatory responsiveness. Using primary human monocytes and peripheral blood mononuclear cells (PBMCs) from healthy donors and sepsis patients, we demonstrate that P7-Pen enhances cytokine production in response to Toll-like receptor stimulation while having minimal effects under basal conditions. P7-Pen treatment increased global histone acetylation, particularly H3 acetylation, and prevented the development of endotoxin tolerance. Transcriptomic and functional analyses revealed restoration of inflammatory gene expression, including *TNF*, *IL6*, and *IFNB1*, in otherwise hyporesponsive cells. Mechanistically, we identified the lysine deacetylase ABHD14B as a direct binding partner of P7-Pen. Silencing of *ABHD14B* recapitulated the effects of P7-Pen, leading to enhanced histone acetylation and cytokine production. Importantly, P7-Pen selectively potentiated responses to TLR ligands without inducing basal hyperinflammation. Collectively, our findings identify ABHD14B-dependent epigenetic regulation as a key checkpoint in innate immune tolerance and establish P7-Pen as a novel tool to restore immune responsiveness in sepsis.

## Introduction

Inflammation plays a crucial role in the development and progression of many diseases [1, 2]. The binding of pathogens or damage-associated molecules to pattern recognition receptors (PRRs), such as Toll-like receptors (TLRs), activates intracellular signaling cascades that initiate host defense responses and induce production of pro-inflammatory cytokines and type I interferons [3]. While essential for pathogen clearance, overstimulation of PRRs can result in uncontrolled systemic or chronic inflammation [4–6]. Therefore, strategies capable of modulating PRR-driven responses are critical for treating inflammatory disorders.

Severe acute inflammatory states are detrimental to host health and may lead to life-threatening complications. Among these, sepsis represents one of the most prevalent and deadly conditions, characterized by rapid onset and high mortality [7]. Globally, sepsis accounts for one in five deaths and kills approximately 11 million people annually, including many children (WHO global report 2020)[8]. Although improved clinical management has reduced short-term mortality, long-term mortality remains alarmingly high—reaching 40–80%—and 10–20% of survivors experience lasting complications such as chronic fatigue, cognitive impairment, and pain[9, 10]. A major contributor to poor outcomes is post-sepsis immunosuppression, which predisposes patients to secondary infections, opportunistic pathogens, and reactivation of latent viruses. The mechanisms underlying this immunosuppressed state are multifactorial and include immune cell depletion or exhaustion and monocyte tolerance [11]. Long-term follow-up studies have shown that monocytes from sepsis survivors exhibit impaired cytokine production upon lipopolysaccharide (LPS) restimulation, a hallmark associated with worse clinical prognosis[9]. Despite substantial advances in inflammation research, no approved therapy exists that attenuates inflammation without simultaneously exacerbating immunosuppression (reviewed in [12]).

Recent evidence suggests that epigenetic mechanisms are extensively perturbed during sepsis progression and may play a pivotal role in the development of immunosuppression at later stages [13, 14]. Epigenetic regulation refers to heritable changes in gene expression that occur without altering the underlying DNA sequence and includes DNA methylation, histone modifications, and non-coding RNA expression[13, 14]. Histone modifications—such as acetylation, methylation, and phosphorylation[14]—are particularly relevant in infection-driven inflammation, with acetylation considered a key regulator of inflammatory responses [15]. Histone acetylation is catalyzed by histone acetyltransferases (HATs), which add acetyl groups to lysines, and histone deacetylases (HDACs), which remove them[14]. Although acetylation was once considered highly unstable, it is now recognized that histone acetylation can persist for days and can function as a platform for recruitment of additional chromatin-modifying factors[16]. Increased histone acetylation generally correlates with enhanced transcription of pro-inflammatory cytokines and antimicrobial mediators[14, 17]. HDACs also regulate numerous non-histone substrates central to metabolism, signaling, and cell cycle control[13]. They are divided into Zn²⁺-dependent classes I, II, IV and NAD⁺-dependent class III sirtuins[18]. Recently, an atypical HDAC—ABHD14B/ABHEB—was identified[19, 20]. Unlike classical HDACs, ABHEB catalyzes deacetylation using CoA to generate acetyl-CoA and free amines [20] and has been implicated in glucose metabolism [19, 20]. Published transcriptomic data indicate that *ABHD14B* expression negatively correlates with hyper-inflammatory signatures in blood cells of sepsis patients during acute shock (GEO repository, dataset GDS427), suggesting a potential anti-inflammatory role[21]. However, its function in inflammation and immunosuppression remains unexplored, and no inhibitors are currently available.

The reversible nature of epigenetic modifications and the druggability of chromatin-modifying enzymes make them attractive therapeutic targets for inflammatory diseases, including sepsis [13, 14, 22, 23]. Yet, the clinical use of HDAC inhibitors remains problematic: reduced HDAC activity does not always correlate with expected transcriptional activation, and several inhibitors have been associated with increased infection susceptibility in cancer patients [24, 25]. Moreover, most available HDAC inhibitors lack specificity and causing considerable off-target toxicity[14, 22, 26]. Among PRRs, TLRs are central to the initiation and propagation of inflammatory responses in sepsis. TLR4 plays a particularly important role as the receptor for LPS, a major component of Gram-negative bacteria, as well as endogenous damage-associated molecules [27]. During severe infection, circulating bacteria or LPS trigger overwhelming inflammation (cytokine storm) and contribute to organ dysfunction. Prolonged or excessive TLR4 signaling promotes immunosuppression [6, 28, 29].

Recently, we developed a cell-penetrating anti-inflammatory peptide, P7-Pen, derived from the signaling lymphocyte activation molecule family member 1 (SLAMF1) [30]. P7-Pen inhibits signaling downstream of TLR4 and TLR7/8 and reduces IFNβ and pro-inflammatory cytokine production in human whole blood and primary immune cells. It selectively suppresses cytokine release by directly targeting the TLR adaptor proteins TRAM and TIRAP [31, 32].

Here, we show that P7-Pen exerts a dual role in the modulation of TLR4-mediated signaling. Transient exposure of immune cells to P7-Pen initially acts as a reversible inhibitor; however, once the peptide is removed, it primes cells for an enhanced response to subsequent TLR4 stimulation. These observations prompted us to investigate the mechanisms underlying this unexpected priming effect. Specifically, we examined whether transient peptide exposure could prevent endotoxin-induced monocyte anergy and enhance TLR4 responsiveness in PBMCs from immunosuppressed sepsis patients. We further found that P7-Pen co-precipitates with the lysine deacetylase ABHD14B and reshapes both the global cellular acetylation profile and histone H3 acetylation patterns. Based on these findings, we propose that ABHD14B is an important regulator of TLR4-driven inflammation and that the peptide’s immunomodulatory activity may be mediated through modulation of ABHD14B function.

## Materials and methods

### Blood withdrawal, sample collection and peripheral blood mononuclear cells (PBMC) isolation and treatment

Experimental procedures with blood samples from healthy donors/volunteers were approved by the Regional Committee for Medical and Health Research Ethics (REK) in Central Norway (no. 2009/2245), and for sepsis patients in potential immunosuppression stage – (no. 2022/ 368316). For the heparin-based blood model we collected blood samples from healthy volunteers from the St. Olavs University Hospital Blood Bank by written informed consent, which was approved (REK#503182). The re-use of these samples as controls for sepsis patient samples from another study was also approved (REK#601801). Blood samples from each participant were collected in 6 mL VACUETTE®NH Sodium Heparin tubes (Greiner Bio-One). PBMC isolation using Lymphoprep (Serumwerk Bernburg, Bernburg, Germany) was performed within 30 min after blood collection. For PBMCs isolation, 15-24 ml of heparinized blood was diluted in PBS by adding 0.5 volume of PBS to the sample to reduce erythrocyte aggregation and increase PBMC yield. Diluted samples were layered on Lymphoprep (1:1 to original sample volume) in 50 ml tubes, and centrifugated for 25 min at 800 g with brakes off. Plasma samples were immediately transferred to PP tubes and stored at −80 prior to further analysis. PBMC were washed 3 times by Hank’s Balanced Salt Solution (HBSS) (Sigma, Merck) containing 2 % FCS and incubated with solvent or peptides for indicated time in RPMI 1640 media (Sigma, Merck) with 5% FCS in 24 well-plates. For media exchange and supernatants collection, plates with PBMC were centrifuged for 10 min at 800 g. Collected supernatants were transferred to PP tubes and stored at −80 prior to further analysis. After treatment and stimulation, PBMC on plates were washed by PBS and lyzed using 0.5 ml of Qiazol lysis buffer (#79306, QIAGEN, Germantown, AR, USA) suitable for simultaneous RNA/protein isolation.

### Heparin-based dilution model

In-house heparin-based dilution model, which minimizes sample handling to reduce experimental bias, was developed based on IFA with whole blood (TruCulture) [10.1016/j.clim.2019.108312] and described in [10.1016/j.mtbio.2025.102113]. Briefly, peripheral venous blood from sepsis patients (n = 5) or healthy donors (n=20) was collected in 6 mL VACUETTE®NH Sodium Heparin tubes (Greiner Bio-One). Within 60 min after blood collection, aliquots of 250 μL were added to polypropylene stimulation tubes (Cluster tubes, Corning) containing 500 μL RPMI cell culture medium, 10 U/mL sodium heparin, and various immune agonists. To minimize technical variation, immune activators were pre-diluted in RPMI in large batches, aliquoted into the stimulation tubes, and stored at −20 ^°^C until use. Once thawed at RT, fresh blood was added at a 1:3 dilution and gently mixed by pipetting 6 times. Samples were incubated at 37 ^°^C and 5 % CO_2_ under static conditions for 18 or 72 h. During incubation, blood cells sedimented via gravity, and supernatants were collected and stored at −20 ^°^C. The stimulant concentration and incubation time were optimised as described [10.1016/j.mtbio.2025.102113]. For 24 h of stimulation, we included the non-stimulated baseline (RPMI alone) and RPMI with selective TLR agonists (Invivogen) – TLR4 (LPS O111:B4, 1 ng/ml), TLR8 (TL8-506, 300 ng/ml) and TLR2 (FSL-1, 100 ng/ml). T cell responses were assessed after 72 h, under non-stimulated conditions (RPMI alone) and RPMI with a synthetic superantigen (CytoStim, 2 μl/ml) from Miltenyi Biotec (cross-binding TCR and MCH) in combination with co-stimulating anti-CD28 mAb (InVivoMAb, BioXCell, 300 ng/ml). Levels of accumulated cytokines in the supernatants were analyzed by Bioplex.

### Cell lines

THP-1 WT (monocytic cell line derived from acute monocytic leukemia ATCC TIB-202) were cultured in RMPI 1640 supplemented by 10% heat-inactivated FCS, 100 U/ml penicillin, 100 μg/ml streptomycin (Thermo Fisher Scientific), and 5 μM β-mercaptoethanol (Sigma-Aldrich, Merck). Prior experimental procedures THP-1 cells were differentiated with 60 ng/ml of phorbol 12-myristate 13-acetate (PMA) (Sigma, Merck) for 48 h, followed by 48 h in a medium without PMA. Media was changed to fresh media before the pretreatments and stimulation. Cell lysates for mass spectrometry were prepared from human multiple myeloma cell line ANBL-6 (from Dr. Diane Jelinek, Mayo Clinic, Rochester, MN, USA). ANBL-6 cells were cultured in RPMI1640 containing 10% FCS (Gibco), pen/strep (Thermo Fisher Scientific) and supplemented with 1 ng/ml rhIL-6.

### Reagents

Synthetic peptides for assays with living cells were from Thermo Fisher Scientific, with sequences RQIKIWFQNRRMKWKK for Penetratin (Pen), ITVYASVTLTG-Pen for P7-Pen, IATYASTALTG-Pen for C3-Pen control peptide. Peptide modifications and characteristics: N-terminal acetylation, C-terminal amidation, >90% purity, guaranteed TFA removal, control for endotoxin levels (less than <10 EU/mg). All peptides’ stock solutions (1 mM) were prepared using sterile distilled water suitable for cell culture (#15230162, Gibco, Fisher Scientific) and stored at −80; working solutions (15 μM) were prepared in cell-culture media immediately before use. Ultrapure K12 LPS (#tlrl-peklps) from *E. coli*, ultrapure thiazoloquinoline compound CL075 (#tlrl-c75), benzazepine analog TL8-506 (#tlrl-tl8506) and FSL-1 TLR2/TLR6 ligand (#tlrl-fsl) were from InvivoGen (San Diego, CA, USA). For stimulation of the primary cells, LPS or FSL-1 were used at concentration 100 ng/mL, TLR7/8 ligands – 1 µg/mL if not indicated otherwise in the figure legends.

### Antibodies

The following primary antibodies were used: mouse β-tubulin (D3U1W, #86298), STAT1 (9H2, #9176), rabbit histone H3 (D1H2, #4499), H3K9ac (acetyl-histone H3 Lys9, C5B11, #9649), H3K27ac (acetyl-histone H3 Lys27, D5E4, #8173), H3K56ac (acetyl-histone H3 Lys56, #4243), H4K5ac (acetyl-histone H4 Lys5, D12B3, #8647), H4K12ac (acetyl-histone H4 Lys12, D2W60, #13944), phospho-STAT1 (Tyr701) (58D6, #9167) from Cell Signaling Technology (Danvers, MA, USA); mouse CD8a (1G2B10, #66868-1-Ig), rabbit CD3 (#17617-1-AP), CD14 (#17000-1-AP) were from Proteintech (); acetylated lysine mouse monoclonal Abs (MA1-2021) were from Invitrogen (Themo Fisher Scientific,); rabbit anti-ABHD14B (in-house, Pune, India) [refs]. Secondary antibodies (HRP-linked) swine anti-rabbit (#P0399) and goat anti-mouse (#P0447) were from DAKO Denmark A/S (Glostrup, Denmark).

### RT-qPCR

Total RNA was isolated from the cells using Qiazol reagent (#79306, QIAGEN, Germantown, AR, USA), and chloroform extraction was followed by purification on RNeasy Mini columns (#74106) with DNAse digestion step (#79254, QIAGEN). cDNA was prepared with a Maxima First Strand cDNA Synthesis Kit for a quantitative real-time polymerase-chain reaction (RT-qPCR) (#K1641, ThermoFisher Scientific, Waltham, MA, USA), in accordance with the protocol of the manufacturer, from 400–600 ng of total RNA per sample. Q-PCR was performed using the PerfeCTa qPCR FastMix (#733-1393, Quanta Biosciences, Gaithersburg, MD, USA) in replicates and cycled in a StepOnePlus™ Real-Time PCR cycler (ThermoFisher Scientific, Waltham, MA, USA). The following TaqMan® Gene Expression Assays (Applied Biosys-tems®, ThermoFisher Scientific) were used: *IFNB1* (Hs01077958_s1), *TNF* (Hs00174128_m1), *TBP* (Hs00427620_m1), *IL-6* (Hs00985639_m1), *IL-1β* (Hs01555410_m1). The level of *TBP* mRNA was used for normalization and the results presented as a relative expression compared to the control’s untreated sample. Relative expression was calculated using Pfaffl’s mathematical model. Graphs and statistical analyses were made with GraphPad Prism v10.1.2 (Dotmatics, Bishops Stortford, UK), with additional details provided in the figure legends or statistics paragraph.

### RNAseq analysis

For the RNAseq analysis, we collected blood samples from healthy volunteers (n=10) at the Rikshospitalet, Oslo University Hospital, after obtaining written informed consent. Blood samples collection and analysis was approved by Regional Ethical Committee in South East Norway. Total RNA was isolated from PBMCs following treatment with vehicle (water) or 15 µM P7-Pen for 16–20 h, as described in the PBMC isolation methods section. RNA quality control, polyA selected short read library preparations were performed at Novogene (UK). Libraries were sequenced on the Illumina platform (Novogene, UK) as 150 bp paired-end reads at depth of >30M reads per sample. Raw reads were pseudoaligned to the Ensemble Release 115 total Human transcriptome (GRCh38) using *kallisto* (v0.56) tool. Per-transcript read count values were aggregated into gene count values using *tximport* Bioconductor library and fed into locally deployed iDEP 2.20 integrated bioinformatic RNAseq analysis pipeline, following filtering of low abundance transcripts (tpm<1 for >3 samples) differentially expressed genes were identified using integrated DESeq2 package. Differentially expressed genes were defined as those with LFC>2 and adjusted p value of <0.1. Pathways were tested with the GSEA (Pre-Ranked) workflow, applying a pathway FDR cutoff of 0.1 and filtering genes with DEG FDR greater than 1.

### ELISA and BioPlex Assays

Cytokine expression for supernatants from healthy donors and sepsis patients PBMCs was analyzed using BioPlex cytokine assays from Bio-Rad, in accordance with the instructions of the manufacturer with the recommended concentration of reagents, but in reduced volume (1:2), using the Bio-Plex Pro™ Reagent Kit III and Bio-Plex™ 200 System (Bio-Rad, Hercules, CA, USA). Human TNF-alpha DuoSet ELISA (#DY210) (R&D Systems, Minneapolis, MN, USA) was used instead of Bio-Plex when indicated in the figure legends.

### Pull-downs by biotinylated peptides

To examine peptide interaction with target proteins, we used biotinylated peptides with biotin covalently linked to C-terminal lysine of Pen and P7-Pen (synthesis and modification done by Thermo Fisher Scientific). For mass spectrometry analysis three independent pull downs (PDs) were performed and analyzed. For each experimental replicate, human ANBL-6 multiple myeloma cells (20 millions) were washed by PBS and lysed in 1X RIPA lysis buffer (150 mM NaCl, 50 mM Tris-HCl, pH 7.5, 1% Triton X-100, 5 mM EDTA, 50 mM NaF, 2 mM Na_3_VO_4_, protease inhibitors (Thermo Fisher Scientific), and phosphatase inhibitors (Roche)) for 30 min at 4 °C with agitation and cleared by centrifugation. Protein concentrations were measured using Pierce™ BCA Protein Assay Kit (Thermo Fisher Scientific) according to the manufacturers protocol, with 500 ug of samples used for PD. PD assays by biotinylated peptides were performed for 30 min at +4 °C with rotation, using Perce MS-compatible Magnetic IP kit (#90408, Thermo Fisher Scientific). Co-precipitated proteins for mass spectrometry analysis were eluted using elution buffer kit and processed for mass spectrometry as described in a separate section. PDs from primary human monocytes for Western blotting (WB) analysis were performed to validate mass spectrometry findings using same experimental conditions except for a different elution step. Co-precipitated proteins for WB were eluted by heating streptavidin beads with 1× loading buffer (Thermo Scientific) containing 40 mM DTT (Merck), followed by SDS-PAGE and WB. Lower parts of gels for PDs were stained by SimplyBlue Safe Stain (Thermo Fisher Scientific) to access peptides’ loading. Images of stained gels were taken on Gel Doc EZ Imager (Bio-Rad, Bio-Rad Laboratories).

### Western Blotting and SDS-PAGE

Cell lysates were prepared by simultaneous extraction of proteins and total RNA using Qiazol reagent (QIAGEN), as suggested by the manufacturer. Protein pellets were dissolved by heating the samples for 10 min at 95 °C in a buffer containing 4 M urea, 1% SDS (#436143, Sigma, Merck), and NuPAGE® LDS Sample Buffer (4X) (#NP0007, Thermo Fisher Scientific), with a final 25 mM DTT in the samples. Otherwise, lysates were made using 1X RIPA lysis buffer with inhibitors of proteases and phosphatases [ref to peptides paper]. Samples were loaded to precast NuPAGE™ Novex™ protein gels (#WG1402BOX, ThermoFisher Scientific). Proteins on gels were transferred to iBlot Transfer Stacks (#IB33001, Thermo Fisher Scientific) by using the iBlot Gel Transfer Device (ThermoFisher Scientific). The blots were developed with the SuperSignal West Femto substrate (#34095, ThermoFisher Scientific) and visualized with the LI-COR ODYSSEY Fc Imaging System (LI-COR Biotechnology, Lincoln, NE, USA). For densitometry analysis of the bands, Odyssey Image Analysis Studio 5.2 software (LI-COR Biotechnology, Lincoln, NE, USA) was used, and the relative numbers of bands’ intensity were normalized to the intensities of the respective loading-control protein (β-tubulin). Loading-control protein expression was always performed on the same membrane as the protein of interest.

### Silencing in primary human monocyte-derived macrophages (MDMs)

Upon isolation of PBMCs, cells were counted using Z2 Coulter particle count and size analyzer (Beckman Coulter) on program B, resuspended in RPMI 1640 (Sigma-Aldrich, Merck) supplemented with 5% of pooled human serum at a concentration of 8 × 10^6^ per ml, and seeded to 24-well (0.5 ml per well) cell culture dishes. After a 45-min incubation allowing surface adherence of monocytes, the dishes were washed three times by HBSS to remove nonadherent cells. MDMs were obtained by differentiating cells for 12 days in RPMI1640 with 10% human serum and 25 ng/mL rhM-CSF (#216-MC-025, R&D Systems, Minneapolis, MN, USA). Silencing of *ABHD14B* gene was performed on day 8 and 9 after isolation using 20 nM silencing oligos (QIAGEN, Germantown, AR, USA) and Lipofectamine 3000 (Invitrogen, ThermoFisher Scientific, Waltham, MA, USA), as suggested by the manufacturer. Oligos used for silencing were AllStars Negative Control siRNA (SI03650318) and FlexiTube Hs_ABHD14B_1 siRNA (SI03192742) – oligo 1, FlexiTube Hs_ABHD14B_2 siRNA (SI04139436) – oligo 2, FlexiTube Hs_MGC15429_2 siRNA (SI00633542) – oligo 3, Hs_MGC15429_3 siRNA (SI00633549) – oligo 4 (QIAGEN).

### Statistical Analysis

Data that were assumed to follow a log-normal distribution was log-transformed prior to statistical analysis. RT-qPCR was log-transformed and analyzed by Repeated Measurements Analysis of Variance (RM-ANOVA), or a mixed model if there was missing data, followed by Holm-Šídák’s multiple comparisons post-test. Bioplex data was analyzed using a Wilcoxon matched-pairs signed-rank test. Data from WB were analyzed using 2-way ANOVA or mixed effect analysis as indicated in the figure legends. All graphs and analyses were generated with GraphPad Prism v10.1.2 (Dotmatics, Bishops Stortford, UK).

## Results

### P7-Pen primes human monocytes and PBMCs for enhanced TLR4 responses and prevents endotoxin-induced anergy

Previously, we showed that P7-Pen peptide significantly decreases TLR4-mediated cytokine mRNA expression and secretion in human monocytes and macrophages, including the THP-1 model system [32]. Based on the chemical properties of peptides—which do not form covalent bonds when interacting with target proteins—we suggested that inhibition of signaling is transient and that the peptide would act as a reversible inhibitor of TLR4-mediated signaling, even after prolonged pre-treatment (16 h). Indeed, peptide pre-treatment significantly decreased LPS-mediated *IFNB1* and *TNF* expression in cells in which the peptide remained present during stimulation (Fig. 1A). However, LPS-mediated cytokine expression was not inhibited in pre-treated cells from parallel wells in which the media were exchanged, and cells were rested for 1 h prior to LPS stimulation (Fig. 1B). Instead, pre-treatment with P7-Pen—but not with the control peptide (Pen)—significantly enhanced LPS-mediated *IFNB1* and *TNF* expression after the peptide was removed from the media (Fig. 1B). Similar results were obtained using primary human peripheral blood mononuclear cells (PBMCs) from healthy donors, which were pre-treated overnight with solvent (water) or peptide, followed by media exchange, 1 h rest, and stimulation with LPS for 2 h (Fig. 1C). Thus, in addition to the reversible inhibition of TLR4-mediated signaling, the peptide primed cells for a more potent response.

**Figure 1.**
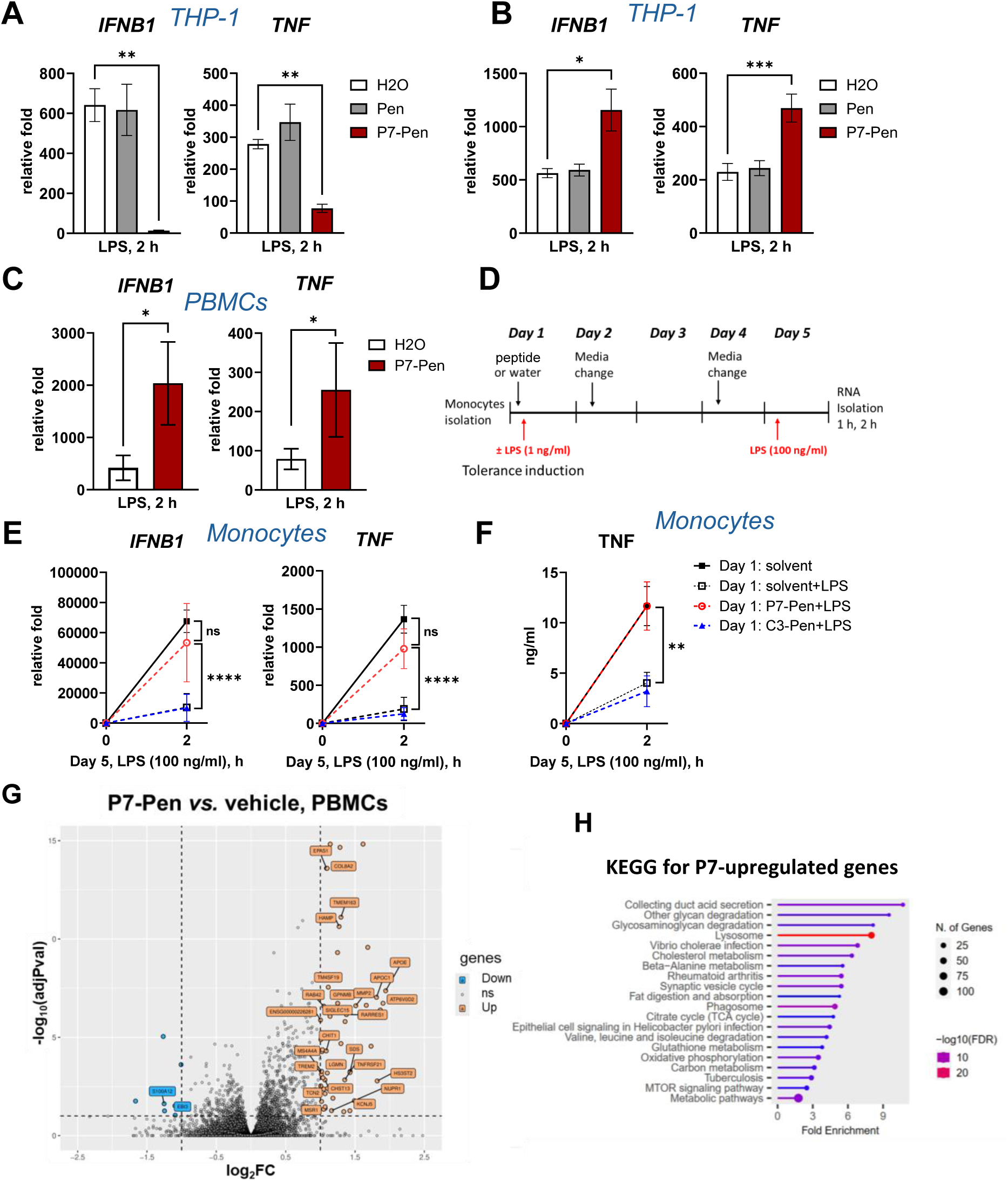
P7-Pen primes monocytes for enhanced LPS responsiveness and prevents tolerance without broadly altering basal transcription. (**A, B**) THP-1 cells were pretreated overnight with vehicle (H_2_O), 15 µM control peptide penetratin (Pen), or 15 µM SLAMF1-derived peptide P7-Pen. Cells were subsequently stimulated with LPS (10 ng/mL) either in the continuous presence of the peptides (**A**) or after media replacement and a 1-h resting period prior to LPS stimulation (**B**). (**C**) Peripheral blood mononuclear cells (PBMCs) isolated from healthy donors were pretreated overnight with vehicle (water) or P7-Pen, followed by media exchange, a 1-h rest period, and stimulation with LPS (100 ng/mL). (**A–C**) Cytokine mRNA expression was quantified by RT-qPCR following LPS stimulation. Data represent mean relative fold change ± SEM from three independent experiments (**A, B**) or from PBMCs of five individual donors (**C**), normalized to unstimulated controls. Statistical significance was assessed by two-way repeated-measures ANOVA on log-transformed data (\**p* < 0.05, \*\**p* < 0.01, \*\*\**p* < 0.001). (**D**) Schematic representation of the experimental model of tolerance induction in primary human monocytes. (**E, F**) RT-qPCR analysis of *IL1B* and *TNF* mRNA expression (**E**) and TNF protein levels measured by ELISA (**F**) on day 5 following tolerance induction. (**G**RNA-seq analysis of PBMCs from ten healthy donors treated with 15 M P7-Pen or vehicle for 16–20 h. P7-Pen induced minor changes in gene expression in PBMCs from a diverse group of healthy individuals. There were 50 genes whose expression significantly (FDR < 0.1) increased more than 2-fold across 10 donor samples with additional 7 genes being downregulated. (**H**) KEGG pathway enrichment analysis of genes upregulated by P7-Pen treatment.

Based on these data (Fig. 1A–C), we hypothesized that the peptide may not only block TLR4-induced cytokine expression during the primary response, as we demonstrated previously [32], but may also prevent the development of monocyte anergy in response to LPS challenge. Monocyte anergy, or endotoxin tolerance, is a hyporesponsive state that develops after an initial exposure to LPS, in which monocytes exhibit markedly reduced production of pro-inflammatory cytokines upon restimulation [33, 34]. Functionally, endotoxin tolerance limits tissue damage during sustained endotoxin exposure (e.g., chronic infection), but it also contributes to sepsis-associated immunoparalysis and heightened susceptibility to secondary infections [33].

Our experimental setup for endotoxin tolerance induction in human monocytes is shown in Fig. 1D. Briefly, monocytes isolated from peripheral blood of healthy donors were either left untreated on day 1 (receiving only solvent control) or stimulated with low-dose LPS (1 ng/ml) on day 1 to induce anergy in media containing solvent (H_2_O), control peptide (C3-Pen), or P7-Pen. On day 5, cells from all conditions were stimulated with LPS (100 ng/ml) for 2 h to assess *IFNB1* and *TNF* expression (Fig. 1E) and TNF secretion (Fig. 1F). Monocytes’ anergy was effectively induced by first LPS challenge (day 1), as shown by reduced cytokine mRNA expression and secretion in response to the second LPS challenge (day 1 solvent *vs.* day 1 solvent + LPS; Fig. 1E, F). Importantly, pre-treatment with P7-Pen, but not with the control peptide C3-Pen or solvent, during the anergy-inducing LPS exposure restored cytokine expression and secretion to levels comparable to non-anergic control (Fig. 1E, F). Thus, P7-Pen prevents anergy development when administered together with the anergy-inducing LPS challenge (Fig. 1D–F).

Hence, we demonstrated that the peptide primed cells for an enhanced response to TLR4 stimulation (Fig. 1A–C). One possible mechanism underlying this priming effect is an altered transcriptional landscape, particularly in genes regulating TLR4 signaling. To investigate this, we assessed the effect of P7-Pen treatment on mRNA expression in primary human PBMCs from healthy donors (n = 10) after 16–20 h by RNA sequencing (Fig. 1G). Peptide treatment exerted only a minor impact on the transcriptome, with 10 significantly downregulated and 68 significantly upregulated genes, none of which were directly associated with regulators of TLR4-mediated pathways (Fig. 1G). KEGG pathway enrichment analysis (ShinyGO) for differentially expressed genes revealed that the peptide-upregulated genes were predominantly enriched in metabolic pathways rather than immune signaling pathways (Fig. 1H). Given the limited transcriptional changes, we considered an alternative mechanism consistent with metabolic reprogramming.

### Protein acetylation profile is modulated by P7-Pen in human PBMCs and monocytes

Protein acetylation—including histone acetylation—is a well-established regulator of cellular metabolism and innate immune training, influencing chromatin accessibility and transcriptional responsiveness without requiring large baseline transcriptional shifts [35–37]. Enhanced histone acetylation has been shown to prime monocytes for augmented cytokine production upon secondary stimulation, in part through epigenetic remodeling of inflammatory loci [38–40]. At the chromatin level, LPS tolerance in monocytes is also associated with epigenetic remodeling, including loss of activating histone acetylation marks and accumulation of repressive marks [41, 42].

Therefore, the modest transcriptional footprint of P7-Pen, together with the metabolic pathway enrichment, led us to hypothesize that peptide-induced priming may be driven by changes in protein acetylation and, specifically, histone acetylation rather than direct transcriptional reprogramming. Histone acetylation is a key chromatin modification that shapes the activity of inflammation-related genes and influences inflammatory responses. All four core histones (H2A, H2B, H3, H4) can be acetylated, though changes on H3 and H4 occur most frequently [43]. Histones acetylation can open up chromatin, allowing transcription factors to bind and activate genes that produce inflammatory mediators [15]. Acetylation marks in histone H3 - H3K9ac, H3K27ac and H3K56ac were shown to be active promoter marks [44–46]], with H3K9ac deacetylation correlating with restrained inflammatory genes expression [47, 48] and TLR stimulation inducing gain of H3K27ac at enhancers near inflammatory genes in macrophages [45]. Acetylation marks in histone H4 - H4K5ac/H4K12ac are “assembly” acetylations on nascent histones, primarily controlled by histone acetyltransferase 1 (HAT1) and linked to replication and global chromatin architecture rather than acute TLR signalling [49, 50].

We therefore examined whether treatment of primary human monocytes and PBMCs with the peptide could alter the total protein acetylation profile (Fig 2A, B) and, in particular, the acetylation of histone H3 at K9, K27, and K56 or H4 at K5 and K12 (Fig. 2).

**Figure 2.**
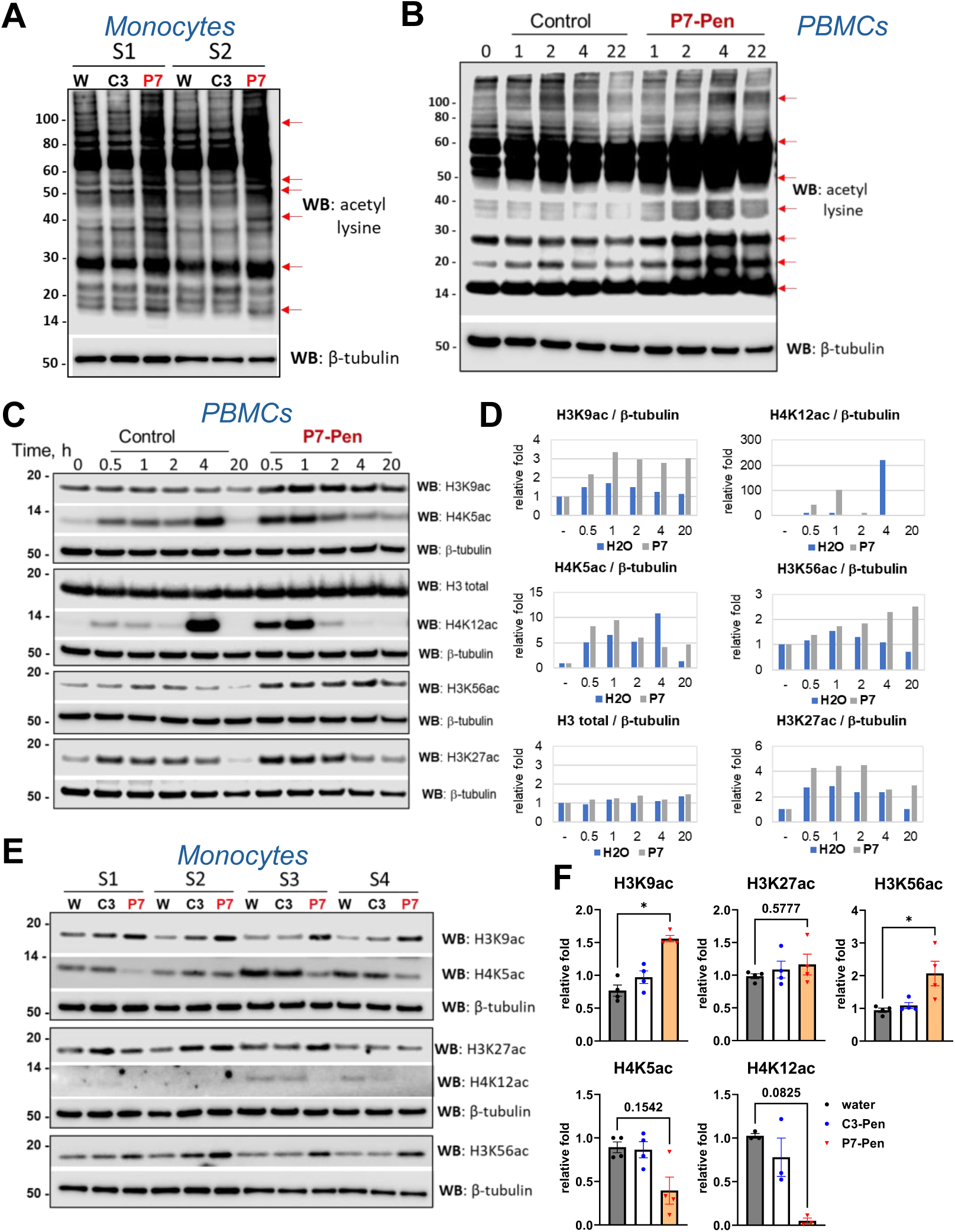
P7-Pen increases global lysine acetylation and histone H3 acetylation in PBMCs and primary monocytes. **(A,B)** Total lysine acetylation was assessed by Western blot in lysates from primary human monocytes **(A)** or PBMCs **(B)** pretreated with vehicle (water), 15 µM control peptide C3-Pen, or 15 µM P7-Pen for 22 h (A) or for the indicated time points (**B**). Representative blots are shown from two of four donors for monocytes (**A**) and from one of three donors for PBMCs (B). Increased abundance of specific acetylated protein bands is indicated by red arrows. β-Tubulin was used as a loading control. **(C)** Representative immunoblots (n = 4) showing histone H3 and H4 acetylation in PBMCs treated with vehicle or 15 µM P7-Pen for the indicated time points. Corresponding densitometric quantification normalized to β-tubulin is shown in **(D)**. **(E)** Western blot analysis of histone H3 and H4 acetylation in primary monocytes (n = 4) treated with vehicle or 15 µM P7-Pen for 22 h. Quantification of histone H3 and H4 acetylation normalized to β-tubulin is shown in **(F)**. Data are presented as mean relative fold change ± SEM. Statistical significance was assessed using nonparametric one-way repeated-measures ANOVA (*p < 0.05*).

Primary human monocytes from healthy donors (n = 4) were incubated for 16 h with peptide solvent (water, W), 15 µM control peptide C3-Pen (C3), or 15 µM SLAMF1-derived peptide P7-Pen (P7). Cell lysates were analyzed by Western blotting using anti-acetyl-lysine antibodies (Fig. 2A). P7-treated cells exhibited a pronounced increase in total protein acetylation across a wide range of molecular weights, indicating broad induction of protein acetylation.

A similar pattern was observed in PBMCs (n = 4), in which P7-Pen induced marked increases in total protein acetylation as early as 2 h after treatment, with elevated acetylation maintained at 4 h and 22 h (Fig. 2B).

Next, we evaluated whether P7-Pen specifically modulates activating histone acetylation marks in PBMCs. Time-course analysis demonstrated that treatment with P7—but not the solvent control—resulted in a strong and rapid increase in H3K9ac, H3K27ac, H3K56ac, as well as H4K5ac and H4K12ac within 30–60 min (Fig. 2C,D; representative donor shown; n = 4). The enhancement of H3 acetylation remained consistent for up to 20 h of treatment. By contrast, H4 acetylation marks (H4K5ac, H4K12ac) displayed greater donor-to-donor variability and fluctuated over time relative to control cells (Fig. 2C,D).

As part of assay validation, we also assessed potential loading controls for normalization of H3 and H4 acetylation signals. Total H3 levels were not altered by P7 treatment and closely overlapped with the H3 signal obtained after probing for acetylation marks (Fig. 2C,D), indicating that H3 itself could in principle serve as an internal control. However, stripping membranes after probing for acetylation marks frequently resulted in incomplete removal of antibodies—particularly for H3 acetylation—which could compromise accurate quantification of total H3 or H4 signals. In contrast, β-tubulin could be reliably detected on the same membranes used for acetylation blots without requiring stripping. Therefore, for consistency and to avoid artifacts from incomplete antibody removal, β-tubulin Western blotting was used for normalization of acetylation signals in all quantifications shown further in the manuscript.

We then examined the acetylation of H3 and H4 in the same monocyte samples used for global acetylation analysis (Fig. 2A; n = 4). Consistent with observations in PBMCs, P7-Pen—but not the control peptide C3—significantly increased H3K9ac and H3K56ac levels after 22 h (Fig. 2A,F). In contrast, H4 acetylation marks (H4K5ac, H4K12ac) were rather reduced in P7-treated monocytes at the 22 h time point compared with solvent or C3-Pen controls (Fig. 2E,F).

Overall, P7-Pen treatment of primary human monocytes and PBMCs induced a rapid and robust increase in acetylation across a broad range of cellular proteins. This included consistent induction of H3K9ac and H3K56ac in monocytes, and H3K9ac, H3K27ac, and H3K56ac in PBMCs. In contrast, H4 acetylation changes were variable and may reflect indirect or secondary regulatory mechanisms. These findings support the idea that P7-Pen induces a chromatin environment conducive to enhanced transcriptional responsiveness without large shifts in baseline gene expression, consistent with a primed or trained-like phenotype.

### Characterization of clinical samples from sepsis patients for potential immunosuppression

Considering the observed effect of transient P7-Pen treatment on PBMCs and monocytes, which resulted in enhanced *IFNB1* and *TNF* expression in response to bacterial endotoxin (Fig. 1C–F), we hypothesized that the peptide could potentially restore immune responsiveness in PBMCs from sepsis patients, particularly by counteracting monocyte anergy. To address this possibility, we collected blood samples from sepsis patients during the early recovery phase (5–8 days after admission to the ICU), with confirmed bacterial infection, along with age- and sex-matched healthy donors. This time window was selected because sepsis-associated immunosuppression typically develops after the initial hyperinflammatory phase and becomes clinically evident between days 3–7 following ICU admission, characterized by reduced cytokine production, monocyte dysfunction, and impaired T-cell responses [12, 51, 52]. Sampling at this stage therefore increases the likelihood of detecting functionally suppressed immune responses relevant to monocyte anergy.

The sepsis cohort comprised ten patients, including equal proportions of Gram-negative and Gram-positive bloodstream infections. Gram-negative pathogens included *Escherichia coli*, *Klebsiella* spp., and *Salmonella* spp., whereas Gram-positive infections were caused by *Staphylococcus aureus*, *Streptococcus dysgalactiae*, and *Streptococcus pneumoniae*. The cohort included 60% female and 40% male patients, with ages ranging from 20 to 80 years (median age 60 years). Patient samples were analyzed in parallel with samples from age- and sex-matched healthy donors, which served as controls.

To determine whether immune suppression was indeed present in this cohort, we first assessed cytokine responsiveness *ex vivo* using a heparinized whole blood stimulation model. The potential immunosuppression in enrolled sepsis patients was evaluated by measuring inflammatory cytokine release in response to TLR agonists—FSL-1/TLR2 (Fig. 3A), LPS/TLR4 (Fig. 3B), and TL8-506/TLR8 (Fig. 3C)—as well as T-cell activation stimuli (synthetic superantigen CytoStim with anti-CD28 mAb) (Fig. 3D). We employed a modified TruCulture-like whole blood system designed to minimize sample handling and reduce experimental bias, originally developed for clinical immune monitoring [53] and later adapted for use in research laboratories [54].

**Figure 3.**
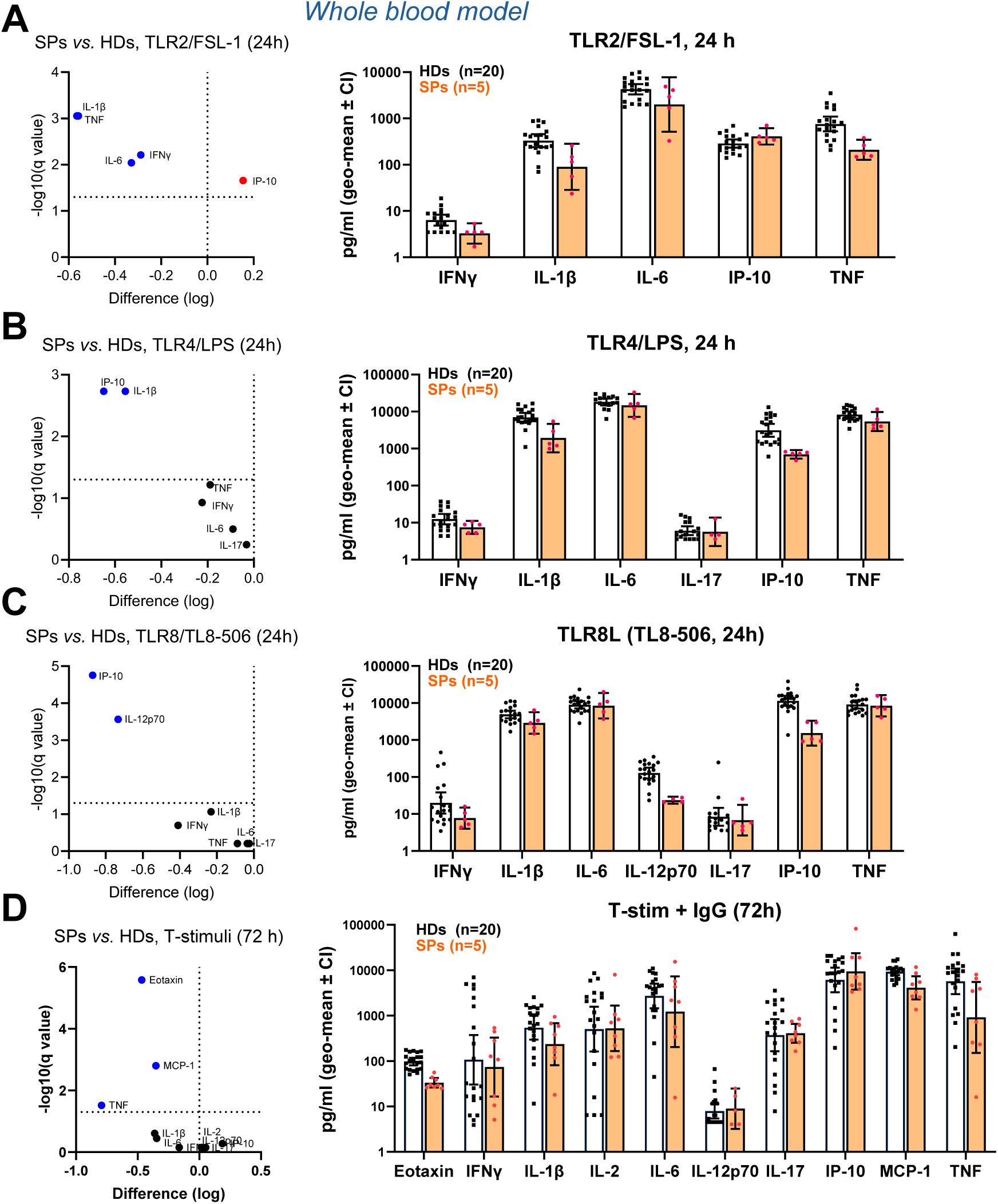
Functional immune profiling reveals immunosuppression in whole blood from sepsis patients. Diluted whole blood samples from sepsis patients (n = 5–8) and healthy donors (n = 20) were stimulated *ex vivo* with defined immune agonists for 24–72 h, and cytokine levels were quantified in diluted plasma supernatants. Volcano plots display log_2_ fold change versus −log_10_ adjusted *p* values (FDR < 0.05). Statistical significance was assessed using multiple unpaired *t*-tests across individual cytokines. **(A)** Stimulation with the TLR2 ligand FSL-1 (100 ng/ml) for 24 h induced significantly attenuated cytokine production in sepsis samples compared with healthy controls, as visualized by a volcano plot and corresponding bar/dot plots. Reduced cytokine levels are shown in blue, while increased IP-10 is shown in red (n = 5). **(B)** Stimulation with the TLR4 ligand LPS (1 ng/ml) for 24 h resulted in significantly reduced production of IP-10 and IL-1β in sepsis samples compared with healthy controls (n = 5). **(C)** Stimulation with the TLR8 ligand TL8-506 (300 ng/ml) for 24 h demonstrated significantly reduced IP-10 and IL-12p70 production in sepsis samples (n = 5). **(D)** Stimulation of diluted whole blood with a T cell/MHC class II–dependent stimulus (Cytostim combined with anti-CD28) for 72 h revealed broadly attenuated cytokine responses in sepsis samples compared with healthy controls (n = 8).

For these experiments, whole blood samples were obtained from a separate cohort of healthy blood donors at the St. Olav’s University Hospital Bloodbank (n = 20, mean age 55 years, 11 females). The whole blood responses of these controls were compared with a subset of the sepsis patients whole blood responses (n = 5 for TLR-ligand stimulation, 4 females, and n=8 for T-cell stimulation, 6 females). The reduced patient number reflects limited blood availability, which was prioritized for other analyses in this study. Cytokine concentrations in plasma after 24 h stimulation were quantified using a Bioplex assay.

Multiple t-test analysis of cytokine release revealed significantly lower secretion of several pro-inflammatory cytokines in sepsis patient whole blood relative to healthy donor controls (q<0.05). Specifically, IL-1β, TNF, IL-6, and IFN-γ responses to TLR2 agonist (Fig. 3A); IP-10 and IL-1β to TLR4 agonist (Fig. 3B); and IP-10 and IL-12p70 response to TLR8 agonist (Fig. 3C) were reduced in sepsis patient samples. Immune responses to T-cell stimulation were also diminished, as indicated by decreased TNF, Eotaxin, and MCP-1 secretion (Fig. 3D). These findings confirmed that patients included in this study exhibited measurable immunosuppression consistent with post-sepsis immune dysfunction.

While highly suitable for evaluating immune responsiveness to TLR ligands and T-cell stimuli, this whole blood assay could not be used to assess the effects of P7-Pen. First, the peptide is likely to be inactivated by heparin present in this model system, as previously demonstrated for other cationic peptides [55]. Consistently, we observed no inhibitory effect of P7-Pen on TLR4-stimulated THP-1 cells when heparin was included at concentrations equivalent to those in whole blood (Supplementary Fig. 1). Heparin alone did not alter LPS-mediated cytokine release; however, it completely abolished the previously established inhibitory effect of P7-Pen [32] on *IFNB1*, *TNF*, *IL6*, and *IL1B* expression in THP-1 cells (Supplementary Fig. 1).

Second, peptide removal from whole blood is not feasible, whereas *in vivo* clearance would normally occur via renal and hepatic filtration. This limitation prevents us from modeling the transient peptide exposure that was critical for the priming effects observed *in vitro* (Fig. 1). Therefore, we proceeded with an alternative experimental setup using isolated PBMCs, as summarized in Fig. 4A.

**Figure 4.**
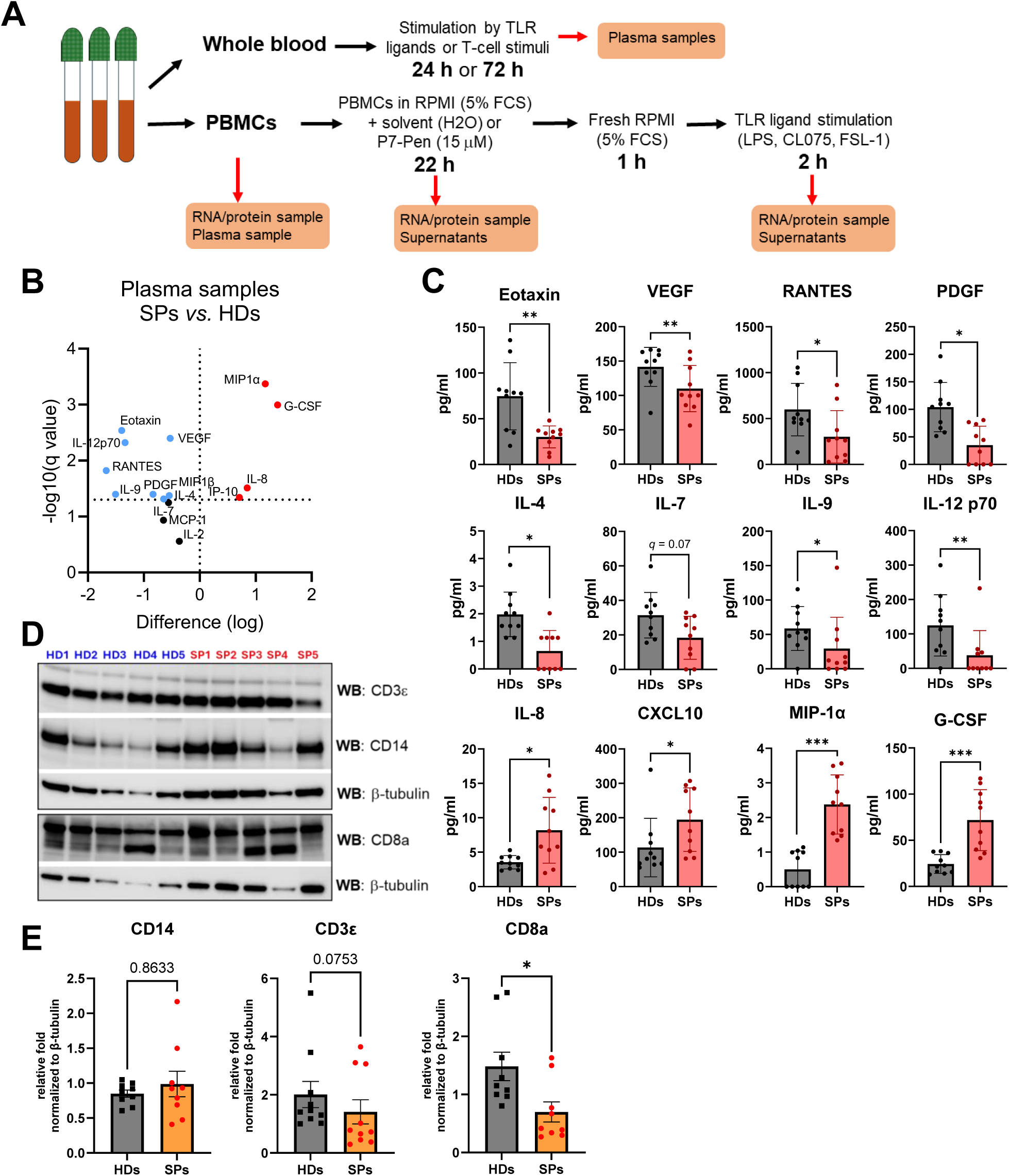
Comparison of cytokine secretion in plasma and immune cell marker expression in PBMCs from sepsis patients and healthy donors. **(A)** Schematic overview of the experimental workflow for blood collection and processing samples from sepsis patients (SPs) and healthy donors (HDs). **(B)** Volcano plots showing cytokine secretion in plasma from SPs versus HDs, generated by plotting log_2_ fold change against −log_10_ FDR-adjusted *p* values (q < 0.05); cytokines decreased in SPs are shown in light blue and increased cytokines in red. **(C)** Quantification of selected cytokines in plasma from HDs and SPs is shown on graphs. Statistical significance for **(B)** and **(C)** was assessed using multiple unpaired *t*-tests (*p* < 0.05, \**p* < 0.01, \*\**p* < 0.001). **(D)** Representative immunoblots of PBMCs from HDs and SPs (n = 10 per group) showing expression of T cell–associated markers (CD3ε, CD8α) and the monocyte marker CD14. **(E)** Densitometric quantification of protein expression normalized to β-tubulin and presented as relative expression levels; statistical significance was assessed using a nonparametric unpaired test (*p* < 0.05).

Analysis of plasma samples from SPs and matched HDs revealed a distinct alteration in circulating cytokine profiles. Compared to HDs, SPs displayed a marked reduction in several cytokines, including Eotaxin, IL-4, IL-9, and RANTES, alongside an increased secretion of pro-inflammatory mediators such as MIP-1α, IP-10, IL-8, and G-CSF (Fig. 4B,C). An overview of the multiplex Bioplex data is presented as a volcano plot with FDR controlled statistics (Fig. 4B), while concentrations of the most strongly altered cytokines are shown individually in Fig. 4C.

Analysis of plasma samples from SPs and matched HDs (characterized in Table 1) revealed a distinct alteration in circulating cytokine profiles. Compared to HDs, SPs displayed a marked reduction in several cytokines, including Eotaxin, IL-4, IL-9, and RANTES, alongside an increased secretion of pro-inflammatory mediators such as MIP-1α, IP-10, IL-8, and G-CSF (Fig. 4B,C). An overview of the multiplex Bioplex data is presented as a multivariate analysis plot (Fig. 4B), while concentrations of the most significantly altered cytokines are shown individually in Fig. 4C.

Similar mixed patterns of cytokine dysregulation have been described in sepsis cohorts, particularly in the early recovery phase (days 3–7). For example, elevated levels of IL-6, IL-8, G-CSF, IP-10, MIP-1α/β and MCP-1 have been reported in sepsis compared with healthy controls and are associated with disease severity and poor outcomes, while significantly increased plasma IL-6, IL-8 and G-CSF were detected in sepsis patients relative to controls [56, 57]. Conversely, reduced levels of chemokines such as RANTES, Eotaxin, and Th2-associated cytokines (e.g., IL-4) have been observed in later or immunosuppressed phases of infection/sepsis, consistent with a transition toward immune exhaustion [56]. These findings align well with our observations of reduced Eotaxin, IL-4, IL-9 and RANTES in SPs, together with persistent elevation of inflammatory mediators.

For further profiling of mononuclear cells in our samples, we analysed the expression of the monocyte marker CD14, the pan-T-cell marker CD3ε, and the cytotoxic T-cell marker CD8α in total protein lysates from PBMCs prepared immediately after isolation, using Western blot (Fig. 4D,E). Figure 4D shows representative blots for these proteins from both HDs and SPs, and quantification normalized to β-tubulin is presented in Figure 4E. Consistent with prior studies of immunosuppressed sepsis patients, the amount of CD14—and thus inferred monocyte number—did not differ significantly between HDs and SPs. This is consistent with prior data in survivors of sepsis, where total monocyte counts either increase or remain stable rather than decrease sharply in the recovery phase [58, 59].

In contrast, there was a clear trend toward reduced CD3ε expression and a statistically significant decrease in CD8α expression in SPs. This reduction of T-cell markers in sepsis aligns with other reports showing lymphopenia and loss of CD8+ T cells in septic patients. For example, Li et al. (2023) documented significantly decreased counts of CD3+ and CD8+ T cells within 24 h of ICU admission in sepsis [60, 61].

Together, these findings confirm that our sepsis patient cohort exhibits a characteristic immunosuppressed phenotype, with preserved monocyte numbers but reduced T-cell markers and impaired cytokine responses. This pattern is typical for the early recovery phase in sepsis survivors, where functional immune defects persist despite partial clinical improvement. Therefore, this cohort provides a clinically relevant immunosuppressed setting and an appropriate experimental model in which to evaluate whether transient P7-Pen treatment can restore immune responsiveness in patient-derived PBMCs.

### Peptide treatment induces cytokine secretion by PBMCs from HDs and SPs

After isolation, PBMCs were diluted in cell culture media containing solvent (water) or 15 µM P7-Pen and incubated for 20–22 h prior to supernatant collection and media exchange (as in setup illustrated in Fig. 4A). Cytokine profiling using a Bioplex assay demonstrated that P7-Pen treatment alone significantly enhanced the release of multiple pro-inflammatory cytokines by PBMCs from HDs compared with solvent-treated controls (Fig. 5A, C). In contrast, PBMCs from SPs displayed a less pronounced increase in cytokine secretion following P7-Pen exposure (Fig. 5B, C). An overview of the multiplex data is shown as differential cytokine expression plots (Fig. 5A, B), while concentrations of the significantly altered cytokines are presented individually in Fig. 5C, allowing direct comparison of cytokine levels released by PBMCs from HDs and SPs. Cytokines that were not significantly affected by P7-Pen treatment are shown in Supplementary Fig. 2. Importantly, baseline cytokine release from solvent-treated PBMCs did not differ significantly between SPs and HDs (Supplementary Fig. 3), indicating that the observed differences were driven by peptide exposure rather than intrinsic baseline secretion in culture.

**Figure 5.**
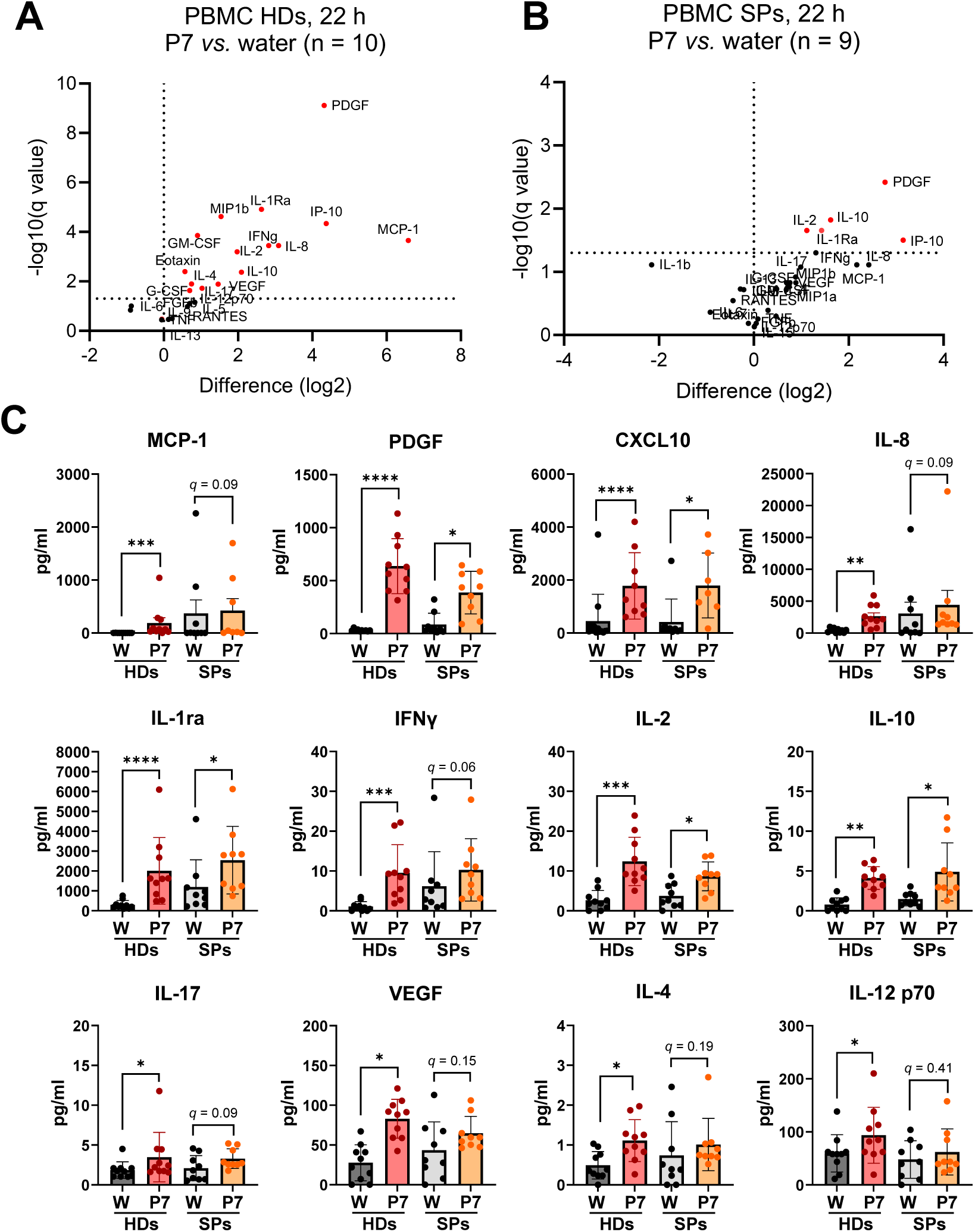
P7-Pen enhances cytokine secretion by PBMCs from healthy donors and sepsis patients. **(A,B)** Volcano plots depicting cytokine secretion by PBMCs treated with vehicle (water) or 15 µM P7-Pen for 20–22 h in samples from healthy donors (A) and sepsis patients (B). Plots were generated by plotting log_2_ fold change against −log_10_ FDR-adjusted *p* values (q < 0.05). **(C)** Quantitative comparison of cytokines significantly upregulated following P7-Pen treatment. Statistical significance for panels **(A–C)** was determined using multiple unpaired *t*-tests (*p* < 0.05, \**p* < 0.01, \*\**p* < 0.001).

Overall, these data show that prolonged P7-Pen treatment in the absence of additional stimulation promoted spontaneous cytokine secretion by PBMCs from HDs, whereas PBMCs from immunosuppressed sepsis patients responded less robustly to the treatment. Notably, this spontaneous cytokine release was consistent with the priming effect observed earlier (Fig. 1) and correlates with the rapid increase in histone H3 acetylation induced by P7-Pen in monocytes and PBMCs (Fig. 2), supporting a mechanism in which peptide-driven chromatin acetylation enhances the transcriptional readiness of immune cells.

### P7-Pen priming enhances TLR4-mediated cytokine responses in PBMCs from both HDs and SPs with less impact on TLR7/8 signaling

Following the 22 h treatment of PBMCs with solvent (water) or P7-Pen, media were exchanged in parallel wells, and cells were rested for 1 h prior to stimulation with LPS (TLR4) or CL075 (TLR7/8) for 2 h. FSL-1 (TLR2) stimulation was performed in a subset of samples and included only in the qPCR analysis. Cytokine responses were assessed by multiplex Bioplex assays (Fig. 6; Supplementary Figs. 4, 6) and RT-qPCR (Supplementary Fig. 5).

**Figure 6.**
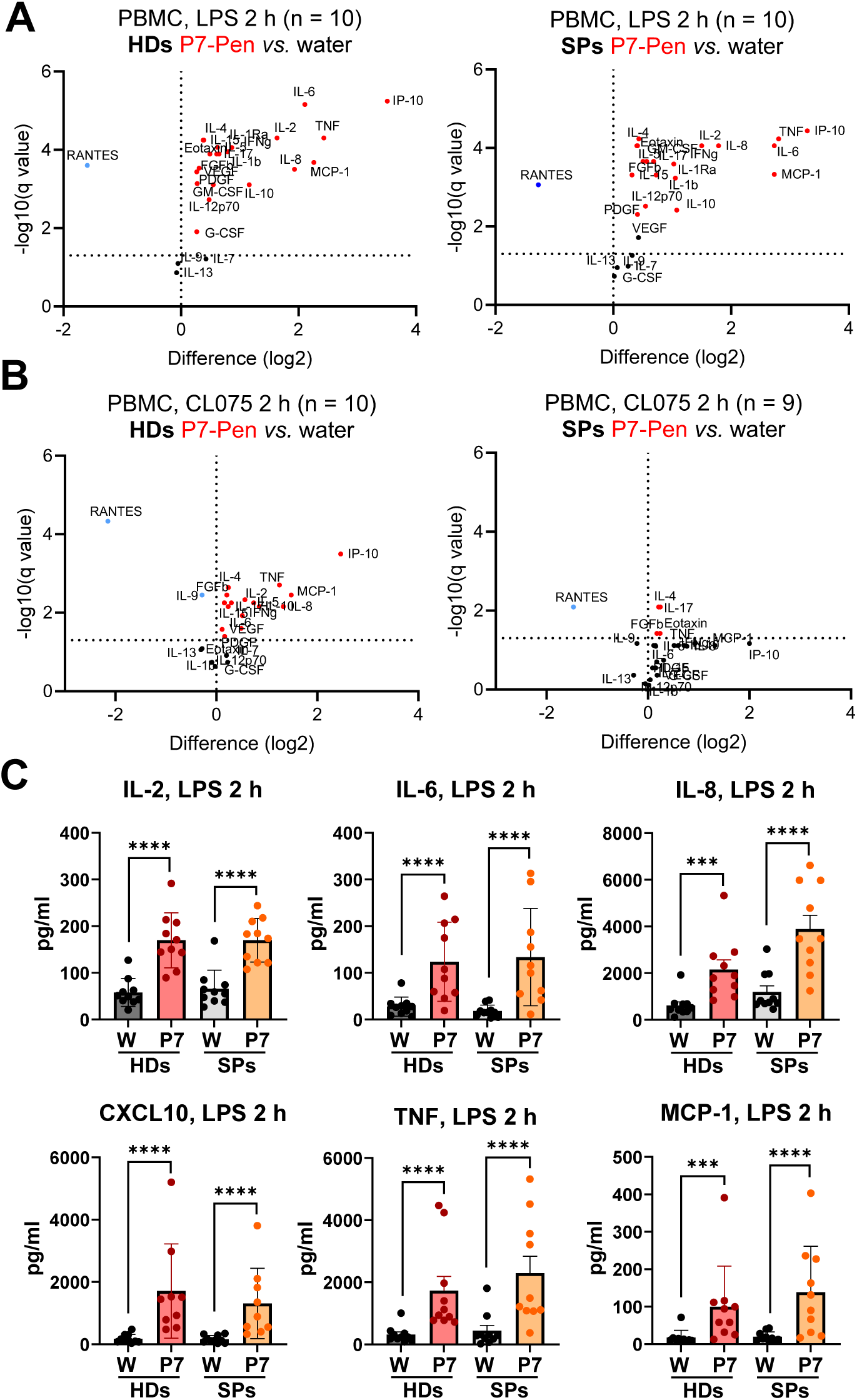
Priming with P7-Pen enhances LPS-induced cytokine secretion in PBMCs from healthy donors and sepsis patients, with minimal effects on CL075-induced responses. Volcano plots comparing cytokine secretion by PBMCs primed with vehicle or P7-Pen and subsequently stimulated with LPS (**A**) or CL075 (**B**) for 2 h. Plots were generated by plotting log_2_ fold change against −log_10_ FDR-adjusted *p* values (q < 0.05). **(C)** Quantification of the most strongly affected LPS-induced cytokines in PBMCs primed with vehicle or P7-Pen. Statistical significance for panels **(A–C)** was assessed using multiple unpaired *t*-tests (*p* < 0.05, \**p* < 0.01, \*\**p* < 0.001).

In solvent-treated controls, *IL6*, *TNF*, and *IL1B* mRNA responses to LPS and CL075 did not differ between PBMCs from HDs and SPs, although LPS-induced *IFNB1* expression was significantly lower in SPs (Supplementary Fig. 4A). Differential analysis indicated that P7-Pen priming significantly enhanced LPS-mediated *IL6*, *IL1B*, *TNF*, and *IFNB1* expression in PBMCs from both HDs and SPs (Supplementary Fig. 4C). In contrast, P7-Pen had minimal effects on CL075 responses: it slightly reduced CL075-induced *IFNB1* and modestly increased CL075-induced *TNF* (<2-fold) in HDs, while showing no effects in SPs (Supplementary Fig. 4D). A trend toward enhanced FSL-1/TLR2-mediated induction of pro-inflammatory cytokines was also observed in P7-Pen–treated PBMCs from both groups (Supplementary Fig. 4E).

At the protein level, Bioplex analysis confirmed the qPCR results. Cytokine secretion from solvent-treated PBMCs did not differ significantly between HDs and SPs for the 2-h LPS or CL075 stimulation (Supplementary Fig. 5A,B). However, P7-Pen priming markedly increased LPS-induced secretion of multiple pro-inflammatory cytokines in both HDs and SPs (Fig. 6A,B). For most cytokines, the fold increase remained below 2-fold, but IL-2, IL-6, IL-8, MCP-1, TNF, and especially IP-10/CXCL10 exhibited more pronounced induction (Figure 6A,B; Supplementary Fig. 5C). Consistent with transcript-level data (Supplementary Fig. 4D), CL075-induced cytokine secretion was minimally affected by P7-Pen in HDs and showed almost no difference compared with solvent controls in SPs (Fig. 6B; Supplementary Fig. 6).

In summary, P7-Pen priming selectively enhanced TLR4-mediated cytokine expression and secretion in PBMCs from both HDs and SPs, while exerting minimal effects on TLR7/8 signaling. This suggests that the priming activity of P7-Pen preferentially targets pathways downstream of TLR4 and possibly TLR2, rather than broadly increasing responsiveness to all TLR ligands.

### Peptide-induced priming correlates with increased H3 acetylation in PBMCs from both HDs and SPs

We next examined whether P7-Pen–mediated priming of PBMCs was associated with changes in histone acetylation. PBMCs from HDs and SPs were treated with solvent, control peptide C3-Pen, or P7-Pen and analyzed for acetylation of histone H3 and H4 lysine residues (Fig. 7; Supplementary Fig. 7). All samples within each group (HDs or SPs) were processed simultaneously to allow direct comparison and statistical analysis, and acetylation levels were normalized to solvent-treated controls.

**Figure 7.**
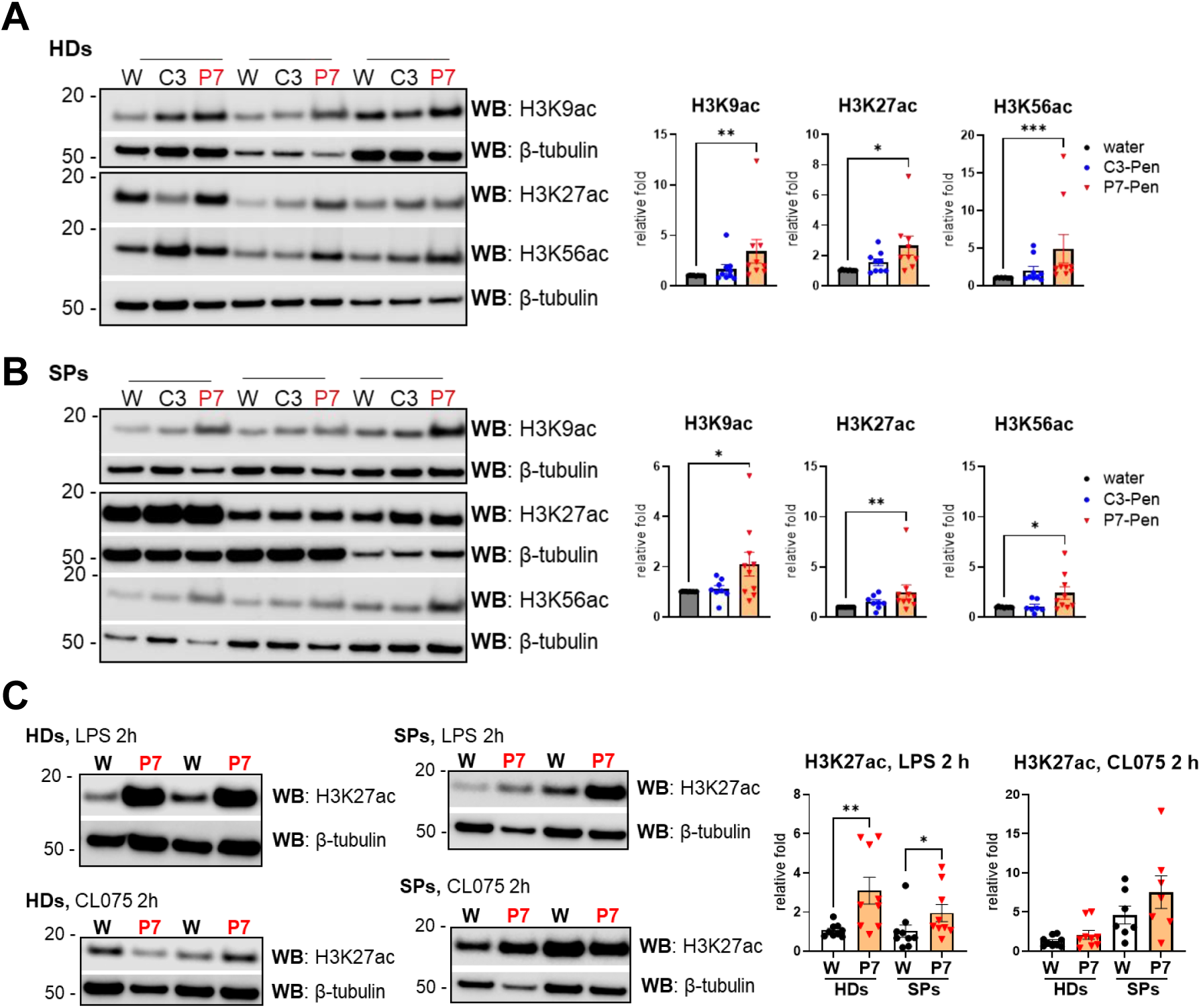
P7-Pen priming increases histone H3 acetylation levels that are maintained during subsequent LPS stimulation of PBMCs. Western blot analysis shows histone H3 acetylation patterns together with quantitative densitometry for all analyzed samples. **(A, B)** Representative immunoblots of PBMC lysates from healthy donors (**A**) or sepsis patients (**B**) showing levels of H3K9ac, H3K27ac, and H3K56ac following priming with vehicle (water), 15 µM control peptide C3-Pen, or 15 µM P7-Pen for 20–22 h. **(C)** Representative immunoblots (2 of 10 donors per group) showing H3K27ac levels in PBMCs primed with water or P7-Pen and subsequently stimulated with LPS or CL075 for 2 h. Quantification shown in the graphs represents signal intensity normalized to the corresponding loading control (β-tubulin). Data are presented as mean relative fold change ± SEM. Statistical significance was assessed using the nonparametric Wilcoxon matched-pairs signed-rank test (*p* < 0.05, \**p* < 0.01, \*\**p* < 0.001).

Consistent with our earlier observations in PBMCs and monocytes (Fig. 2C–F), P7-Pen treatment significantly increased H3 acetylation at K9, K27, and K56 in PBMCs from both HDs and SPs, whereas the control peptide C3-Pen had no significant effect (Fig. 7A,B). The increased H3 acetylation correlated with the enhanced spontaneous cytokine secretion observed after 22 h of peptide treatment (Fig. 5), supporting a link between P7-Pen–induced chromatin acetylation and priming of inflammatory responses.

Due to limited sample availability, downstream analyses after TLR stimulation focused on H3K27ac, a histone mark strongly implicated in long-lasting epigenetic regulation of inflammatory tolerance. Notably, persistent loss of H3K27ac has been identified as a defining chromatin feature of endotoxin- or toxin-induced tolerance, including a study showing that anthrax lethal toxin drives stable H3K27 deacetylation [62]. In P7-Pen–primed PBMCs, LPS stimulation resulted in significantly higher H3K27ac levels compared with solvent controls in both HDs and SPs, whereas CL075 stimulation did not alter H3K27ac (Fig. 7C). This pattern aligns with functional data showing selective enhancement of TLR4-, but not TLR7/8-mediated, cytokine responses in P7-Pen–treated PBMCs (Fig. 6).

In contrast to H3 acetylation, P7-Pen did not significantly alter H4 acetylation at K5 or K12 in either group, whether assessed after 22 h peptide treatment (Supplementary Fig. 7A,B) or subsequent LPS or CL075 stimulation (Supplementary Fig. 7C). These results are consistent with our earlier observations in monocytes and PBMCs (Fig. 2C-F) and indicate that H4 acetylation marks are not substantially affected by P7-Pen in this experimental setting.

In conclusion, P7-Pen induces a robust and selective increase in activating H3 acetylation marks—both at baseline and during TLR4 stimulation—in PBMCs from HDs and SPs. This histone signature parallels the peptide’s priming effect on cytokine production and supports a model in which P7-Pen enhances TLR4 responsiveness through targeted chromatin remodeling at H3-associated regulatory sites.

### Potential mechanism: P7-Pen peptide interacts with the atypical lysine deacetylase ABHD14B and functionally mimics its silencing

To explore the mechanism underlying P7-Pen–induced increases in histone H3 acetylation, we next examined whether the peptide interacts with histone acetyltransferases (HATs) or histone deacetylases (HDACs), which either deposit acetyl groups on lysine residues or remove them respectively. As part of an independent study investigating P7-Pen in cancer cells, we previously performed a mass spectrometry–based pull-down screen in the ANBL6 multiple myeloma cell line to identify proteins co-precipitating with biotinylated P7-Pen compared with control Pen peptide. Re-analysis of these data for potential interactions with chromatin-modifying enzymes revealed a single notable hit: the unconventional lysine deacetylase ABHD14B (also known as ABHEB). Canonical HDACs, including HDAC1, SIRT1, and SIRT3—displayed very low scores and low probabilities of interaction (Fig. 8A). To validate this interaction, we performed pull-down assays using lysates from primary human macrophages and biotinylated peptides. Western blot analysis confirmed that ABHD14B co-precipitated with P7-Pen, but not with the control Pen peptide (Fig. 8B). TRAM, a previously validated P7-Pen target [32], was included as a positive control and was also enriched in P7-Pen pull-downs.

**Figure 8.**
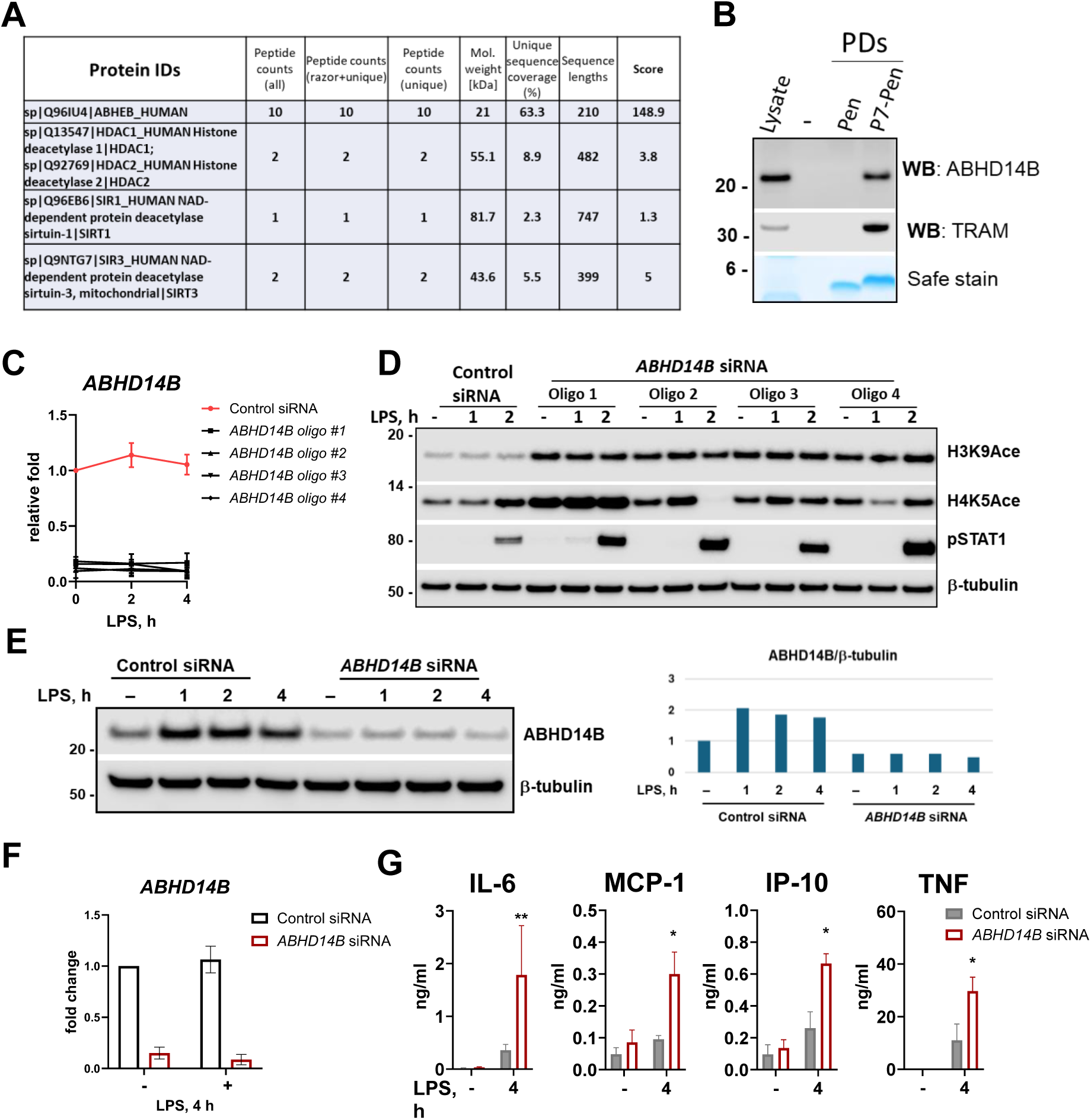
P7-Pen associates with the lysine deacetylase ABHD14B, whose expression inversely correlates with LPS-induced cytokine production. **(A)** Table shows results of a mass spectrometry–based screen identifying lysine deacetylases co-precipitating with P7-Pen compared with control peptide (Pen), based on three independent experiments. **(B)** Western blot analysis of ABHD14B and TRAM (known P7-interacting protein, positive control) in pulldown samples from monocyte lysates using biotinylated Pen or P7-Pen peptides immobilized on streptavidin beads, with whole-cell lysates included as input controls (7.5% of total protein); total protein staining by SimplyBlue is shown as a loading control for peptides. **(C)** The efficiency of *ABHD14B* silencing in primary human macrophages was assessed by RT–qPCR following transfection with four independent A*BHD14B*-targeting siRNAs, with results expressed as mean relative fold change ± SEM from three independent experiments. **(D)** Representative immunoblots illustrate the effects of *ABHD14B* knockdown on histone acetylation (H3K9ac and H4K5ac) and STAT1 phosphorylation, with β-tubulin used as a loading control. **(E–G)** Functional validation of *ABHD14B* silencing was performed in macrophages from six healthy donors, demonstrating efficient knockdown at the protein level (**E**), reduced transcript abundance by qPCR (**F**), and altered LPS-induced secretion of IL-6, MCP-1, IP-10, and TNF (**G**) as measured by multiplex cytokine analysis. Data are presented as mean relative fold change ± SEM, and statistical significance was determined using nonparametric one-way repeated-measures ANOVA (*p* < 0.05).

ABHD14B is structurally and functionally distinct from classical HDACs, exhibiting unique catalytic features and substrate specificity, and its role in inflammation remains largely uncharacterized. To assess whether reduced ABHD14B expression alters the acetylation landscape or inflammatory signaling, we silenced *ABHD14B* in primary monocyte-derived macrophages (MDMs) using four independent siRNA oligos (Fig. 8C–G). All oligos decreased *ABHD14B* mRNA levels by at least 70% (Fig. 8C). ABHD14B knockdown consistently increased H3K9 acetylation and enhanced STAT1 phosphorylation (Fig. 8D). The latter could serve as a surrogate for increased type I interferon signaling in human macrophages as we have shown before [31, 32]. In contrast, effects on H4K5 acetylation were variable across oligos, similar to the minimal impact of P7-Pen on H4 acetylation observed earlier (Fig. 2E,F).

Further, *ABHD14B* siRNA oligo 2 was used for screening the effect of knocking down *ABHD14B* expression in MDMs from several donors to tack the effect of silencing on LPS-induced cytokine secretion (Fig. 8E). Indeed, reduced expression of ABHD14B on protein and mRNA levels (Fig. 8E,G) resulted in the enhanced secretion of pro-inflammatory cytokines IL-6, MCP-1m IP-10 and small increase in TNF secretion in response to LPS stimulation (Fig 8E). Using the siRNA oligo #2, which showed the most consistent silencing efficiency across donor cells, we further evaluated cytokine secretion in response to LPS in MDMs from multiple donors. Reduced ABHD14B expression at both protein and mRNA levels (Fig. 8E,G) resulted in enhanced secretion of IL-6, MCP-1, IP-10, and TNF following LPS stimulation (Fig. 8E). These findings demonstrate for the first time that ABHD14B acts as a negative regulator of pro-inflammatory signaling in human macrophages and that its loss promotes increased H3K9ac and heightened responsiveness to TLR4 stimulation.

Given that P7-Pen directly co-precipitates with ABHD14B and that both P7-Pen treatment and *ABHD14B* knockdown similarly enhance H3 acetylation and amplify LPS-driven cytokine responses, we considered whether the peptide modulates ABHD14B function. Using a gel-based activity-based protein profiling assay with recombinant wild-type and catalytically inactive ABHD14B [20, 63], we found no evidence that P7-Pen (4–20 μM) inhibits ABHD14B enzymatic activity in vitro (Supplementary Fig. 8A–C). Furthermore, although ABHD14B expression fluctuated naturally during PBMC culture, P7-Pen did not alter its expression compared with solvent controls at most time points examined (Supplementary Fig. 8D).

Thus, while P7-Pen interacts with ABHD14B and phenocopies key outcomes of *ABHD14B* silencing—including increased H3K9ac and enhanced TLR4 responsiveness—our data indicate that the peptide does not directly inhibit ABHD14B catalytic activity or expression. Additional work is required to determine whether P7-Pen affects ABHD14B stability, post-translational modification, subcellular localization, or interaction with protein partners to modulate its functional output.

## Discussion

In this study, we identify the SLAMF1-derived peptide P7-Pen as a reversible modulator of TLR4 signaling that could both inhibit acute cytokine responses and prime human monocytes and PBMCs for enhanced reactivity to subsequent endotoxin challenge. In line with our previous work [32], P7-Pen acutely suppressed LPS-induced *IFNB1* and *TNF* expression when present during stimulation. However, when the peptide was removed prior to LPS exposure, pre-treated monocytes and PBMCs from healthy donors exhibited significantly increased *IFNB1* and *TNF* induction compared with solvent or control peptides, demonstrating a primed state. Importantly, in *in vitro* model of endotoxin tolerance, P7-Pen prevented the development of monocyte anergy when present during the initial low-dose LPS exposure. Peptide restored cytokine expression and secretion to levels comparable to non-tolerized cells upon secondary LPS stimulation. These data indicate that P7-Pen can both transiently inhibit TLR4 signaling and, on a longer timescale, reprogram monocytes toward heightened responsiveness to subsequent endotoxin exposure.

RNA-sequencing of PBMCs treated with P7-Pen revealed only modest changes in steady-state gene expression, with a limited set of differentially expressed genes and no clear enrichment for canonical regulators of TLR4 signaling. Instead, KEGG analysis showed a predominance of metabolic pathways among upregulated genes. This modest transcriptional footprint, combined with the pronounced functional priming phenotype, argues against a mechanism relying solely on direct transcriptional rewiring at baseline. Rather, our data are consistent with a model in which P7-Pen induces an epigenetic and metabolic state that “licenses” a more robust transcriptional response upon subsequent TLR4 engagement. Such a configuration resembles features of trained immunity, where innate cells acquire enhanced responsiveness through epigenetic and metabolic reprogramming without large changes in basal transcript levels [35–37].

We therefore focused on protein acetylation, a key link between metabolism and chromatin regulation. P7-Pen rapidly and robustly increased total protein acetylation in both monocytes and PBMCs. At the chromatin level, P7-Pen consistently enhanced acetylation of histone H3 at K9, K27, and K56 in PBMCs and monocytes, whereas acetylation of histone H4 at K5 and K12 was more variable and at later time points even reduced. H3K9ac, H3K27ac, and H3K56ac are well-established activating marks associated with open chromatin and inducible gene expression [44–46], and reductions in H3K9ac or H3K27ac have been linked to suppression of inflammatory genes and tolerance [41, 42, 47, 48]. In contrast, H4K5ac/H4K12ac are linked to replication-coupled chromatin assembly and higher-order architecture rather than acute TLR signaling [49, 50]. The preferential and sustained induction of H3K9ac/H3K27ac/H3K56ac, with limited and inconsistent changes in H4K5ac/H4K12ac, thus supports selective engagement of regulatory H3-associated loci rather than a global increase in acetylation.

Functionally, this H3-centered acetylation pattern was tightly linked to P7-Pen–induced priming. Prolonged P7-Pen treatment alone promoted spontaneous secretion of multiple pro-inflammatory cytokines by PBMCs from healthy donors, with PBMCs from sepsis patients showing a less robust response. Following peptide removal from media, P7-Pen priming selectively enhanced TLR4-mediated *IL6*, *IL1B*, *TNF*, and *IFNB1* expression and cytokine secretion in PBMCs from both healthy donors and sepsis patients, while exerting minimal effects on TLR7/8 responses. Consistent with this functional selectivity, P7-Pen–primed PBMCs displayed increased H3K27ac upon LPS stimulation, but not following CL075 stimulation, and H4K5ac/H4K12ac remained largely unchanged. Together, these observations support a model in which P7-Pen enhances TLR4 responsiveness through targeted chromatin remodeling at H3-associated regulatory regions, thereby lowering the threshold for gene activation in response to endotoxin without broadly amplifying responses to all TLR ligands.

Further we explored the relevance of these findings in a disease-relevant immune state by studying PBMCs from sepsis patients in the early recovery phase (5–7 days after ICU admission), a time window associated with the onset of sepsis-induced immunosuppression [12, 51, 52]. Whole blood assays revealed reduced TLR2-, TLR4-, and TLR8-mediated cytokine release and diminished T-cell–dependent responses in sepsis patients compared with healthy donors, consistent with post-sepsis immune dysfunction. Plasma cytokine profiling showed a mixed pattern of persistent inflammation (e.g., elevated IP-10, IL-8, G-CSF, MIP-1α) together with reduced levels of Eotaxin, IL-4, IL-9, and RANTES, in line with previously described dysregulated cytokine signatures in the early recovery phase of sepsis [56, 57]. At the cellular level, PBMCs from sepsis patients exhibited preserved CD14 expression but reduced CD3ε and CD8α, indicating relative maintenance of monocyte numbers along with T-cell depletion, a hallmark of sepsis-associated immunosuppression [58–60]. Within this immunosuppressed yet clinically stable survivor cohort, transient P7-Pen priming still enhanced LPS-induced cytokine production and increased activating H3 acetylation marks, demonstrating that epigenetic priming by P7-Pen can operate in a clinically relevant sepsis setting.

Mechanistically, we identify the atypical lysine deacetylase ABHD14B/ABHEB as a candidate mediator linking P7-Pen to altered H3 acetylation and TLR4 priming. Mass spectrometry–based pull-down analysis in myeloma cells and subsequent validation in primary human monocytes showed that P7-Pen, but not control peptide, co-precipitated with ABHD14B, whereas classical HDACs and sirtuins showed low interaction scores. Silencing ABHD14B in monocyte-derived macrophages increased H3K9ac and enhanced STAT1 phosphorylation and LPS-induced secretion of IL-6, MCP-1, IP-10, and (modestly) TNF, phenocopying key aspects of P7-Pen treatment. These findings identify ABHD14B as a negative regulator of H3 acetylation and TLR4-driven cytokine responses in human macrophages and suggest that P7-Pen peptide may exert its priming effect, at least in part, by functionally modulating ABHD14B.

However, our data argue against direct catalytic inhibition or transcriptional repression of ABHD14B by P7-Pen. In cell-free activity-based protein profiling assays using recombinant ABHD14B, P7-Pen did not inhibit enzyme activity. Moreover, although ABHD14B expression fluctuated during PBMC culture, P7-Pen did not consistently alter its expression compared with controls. These observations suggest that P7-Pen may influence ABHD14B function through alternative mechanisms, such as altering its post-translational modification, subcellular localization, or interaction with protein partners. Future work will be required to dissect how P7-Pen binding to ABHD14B translates into increased H3 acetylation at inflammatory loci and whether this involves re-routing ABHD14B away from chromatin-associated substrates or interfering with its recruitment to specific complexes.

Histone acetylation is tightly linked to cellular metabolic state through the availability of acetyl-CoA, NAD⁺, and other cofactors that regulate HAT and HDAC activities [64, 65]. Monocyte tolerance and sepsis-associated immunosuppression are characterized by metabolic rewiring toward fatty acid oxidation, mitochondrial dysfunction, and impaired glycolytic flux—changes that reduce histone acetylation and restrict chromatin accessibility at pro-inflammatory loci [66,67]. The reduction of H3K27ac and H3K9ac at TLR-responsive enhancers is a hallmark of tolerant monocytes and correlates with diminished TNF and IFN-β production [68–70]. In this context, the ability of P7-Pen to restore H3 acetylation and overcome endotoxin-induced anergy suggests that the peptide may counteract metabolic constraints that enforce epigenetic silencing during tolerance.

Despite some important findings, our study has limitations. The sample size for sepsis patients was small and restricted to survivors with limited organ dysfunction, which may not capture the full spectrum of sepsis-induced immunosuppression. All functional experiments were performed *ex vivo*, and pharmacokinetic factors such as peptide stability, biodistribution, and clearance—as well as potential off-target effects—remain to be addressed *in vivo*. Furthermore, while we demonstrate robust correlations between P7-Pen treatment, increased H3 acetylation, *ABHD14B* silencing, and enhanced TLR4 responses, definitive proof that ABHD14B is the central mediator of P7-Pen’s epigenetic effects will require complementary genetic (e.g., CRISPR knockout or rescue) and biochemical approaches, as well as genome-wide mapping of chromatin changes (e.g., H3K9ac/H3K27ac ChIP-seq) at P7-Pen–responsive loci. Despite these limitations, our findings provide a conceptual framework for how a reversible peptide inhibitor of TLR4 can reprogram innate immune cells toward enhanced responsiveness rather than tolerance. P7-Pen’s ability to raise activating H3 acetylation marks and specifically enhance TLR4/TLR2 signals—while sparing TLR7/8 responses—illustrates how fine-tuned epigenetic adjustments can determine whether immune cells remain unresponsive or shift toward a more alert, primed functional state.

From a translational perspective, such an approach could be exploited to boost antibacterial defenses in patients with sepsis-associated immunosuppression or to enhance vaccine adjuvant efficacy, provided that the risk of excessive inflammation is carefully managed. Future studies should investigate the genome-wide distribution of P7-Pen–induced H3 acetylation, define the interactome of ABHD14B in myeloid cells, and explore whether transient peptide-based modulation of acetylation can be harnessed safely *in vivo* to restore immune competence in sepsis and other states of innate immune paralysis.

Overall, this work lays the foundation for developing peptide-based strategies to restore immune function in conditions marked by impaired monocyte activation.

## Author contributions

SU: formal analysis, investigation, writing - editing; BE: methodology, investigation, resources, writing - editing; JS: methodology, investigation, writing – editing; SP, YS, IBM, VB, IBM, MD, SSK, LR, HV, TBD: investigation, writing – editing; BEH, JKD, TE: methodology, resources, writing - editing; MY - conceptualization, resources, formal analysis, supervision, funding acquisition, investigation, methodology, project administration, and writing - original draft, review, and editing.

## Data availability

All data supporting the findings of this study are available within the article and its supplementary materials. Questions regarding additional methodological details or materials are welcome and can be directed to the corresponding author.

## References

1. Chavda, V.P., J. Feehan, and V. Apostolopoulos, Inflammation: The Cause of All Diseases. Cells, 2024. 13(22).

2. Chen, L., et al., Inflammatory responses and inflammation-associated diseases in organs. Oncotarget, 2018. 9(6): p. 7204–7218.

3. Fitzgerald, K.A. and J.C. Kagan, Toll-like Receptors and the Control of Immunity. Cell, 2020. 180(6): p. 1044–1066.

4. Chen, R., et al., Pattern recognition receptors: function, regulation and therapeutic potential. Signal Transduct Target Ther, 2025. 10(1): p. 216.

5. Kumar, V., Toll-like receptors in sepsis-associated cytokine storm and their endogenous negative regulators as future immunomodulatory targets. Int Immunopharmacol, 2020. 89(Pt B): p. 107087.

6. Wang, K., et al., Toll-like receptors in health and disease. MedComm (2020), 2024. 5(5): p. e549.

7. Hotchkiss, R.S., et al., Sepsis and septic shock. Nat Rev Dis Primers, 2016. 2: p. 16045.

8. Rudd, K.E., et al., *Global, regional, and national sepsis incidence and mortality*, *1990-2017: analysis for the Global Burden of Disease Study*. Lancet, 2020. 395(10219): p. 200–211.

9. Delano, M.J. and P.A. Ward, The immune system’s role in sepsis progression, resolution, and long-term outcome. Immunol Rev, 2016. 274(1): p. 330–353.

10. Mostel, Z., et al., Post-sepsis syndrome - an evolving entity that afflicts survivors of sepsis. Mol Med, 2019. 26(1): p. 6.

11. Biswas, S.K. and E. Lopez-Collazo, Endotoxin tolerance: new mechanisms, molecules and clinical significance. Trends Immunol, 2009. 30(10): p. 475–87.

12. Padovani, C.M. and K. Yin, Immunosuppression in Sepsis: Biomarkers and Specialized Pro-Resolving Mediators. Biomedicines, 2024. 12(1).

13. Cross, D., et al., Epigenetics in Sepsis: Understanding Its Role in Endothelial Dysfunction, Immunosuppression, and Potential Therapeutics. Front Immunol, 2019. 10: p. 1363.

14. Wu, D., et al., Epigenetic mechanisms of Immune remodeling in sepsis: targeting histone modification. Cell Death Dis, 2023. 14(2): p. 112.

15. Gao, S., et al., Histone acetylation: A key regulator of inflammatory responses. Life Sci, 2025. 380: p. 123936.

16. Pereira, J.M., M.A. Hamon, and P. Cossart, A Lasting Impression: Epigenetic Memory of Bacterial Infections? Cell Host Microbe, 2016. 19(5): p. 579–82.

17. Barnes, P.J., I.M. Adcock, and K. Ito, Histone acetylation and deacetylation: importance in inflammatory lung diseases. Eur Respir J, 2005. 25(3): p. 552–63.

18. Park, S.Y. and J.S. Kim, A short guide to histone deacetylases including recent progress on class II enzymes. Exp Mol Med, 2020. 52(2): p. 204–212.

19. Rajendran, A., et al., A multi-omics analysis reveals that the lysine deacetylase ABHD14B influences glucose metabolism in mammals. J Biol Chem, 2022. 298(7): p. 102128.

20. Rajendran, A., et al., Functional Annotation of ABHD14B, an Orphan Serine Hydrolase Enzyme. Biochemistry, 2020. 59(2): p. 183–196.

21. Wynn, J.L., et al., The influence of developmental age on the early transcriptomic response of children with septic shock. Mol Med, 2011. 17(11-12): p. 1146–56.

22. Zhang, S.Y., et al., Histone deacetylases and their inhibitors in inflammatory diseases. Biomed Pharmacother, 2024. 179: p. 117295.

23. You, J., Y. Li, and W. Chong, The role and therapeutic potential of SIRTs in sepsis. Front Immunol, 2024. 15: p. 1394925.

24. Galli, M., et al., A phase II multiple dose clinical trial of histone deacetylase inhibitor ITF2357 in patients with relapsed or progressive multiple myeloma. Ann Hematol, 2010. 89(2): p. 185–90.

25. Ryan, Q.C., et al., Phase I and pharmacokinetic study of MS-275, a histone deacetylase inhibitor, in patients with advanced and refractory solid tumors or lymphoma. J Clin Oncol, 2005. 23(17): p. 3912–22.

26. von Knethen, A. and B. Brune, Histone Deacetylation Inhibitors as Therapy Concept in Sepsis. Int J Mol Sci, 2019. 20(2).

27. Tsujimoto, H., et al., Role of Toll-like receptors in the development of sepsis. Shock, 2008. 29(3): p. 315–21.

28. Cohen, J., The immunopathogenesis of sepsis. Nature, 2002. 420(6917): p. 885–91.

29. Rackov, G., et al., The Role of IFN-beta during the Course of Sepsis Progression and Its Therapeutic Potential. Front Immunol, 2017. 8: p. 493.

30. Nilsen, K.E., et al., TIRAP/Mal Positively Regulates TLR8-Mediated Signaling via IRF5 in Human Cells. Biomedicines, 2022. 10(7).

31. Nilsen, K.E., et al., A novel TIRAP-MyD88 inhibitor blocks TLR7- and TLR8-induced type I IFN responses. J Immunol, 2025.

32. Nilsen, K.E., et al., Peptide derived from SLAMF1 prevents TLR4-mediated inflammation in vitro and in vivo. Life Science Alliance, 2023. 6(12): p. e202302164.

33. Lopez-Collazo, E. and C. Del Fresno, Endotoxin tolerance and trained immunity: breaking down immunological memory barriers. Front Immunol, 2024. 15: p. 1393283.

34. Seeley, J.J. and S. Ghosh, Molecular mechanisms of innate memory and tolerance to LPS. J Leukoc Biol, 2017. 101(1): p. 107–119.

35. Cameron, A.M., S.J. Lawless, and E.J. Pearce, Metabolism and acetylation in innate immune cell function and fate. Semin Immunol, 2016. 28(5): p. 408–416.

36. Ferreira, A.V., et al., Metabolic Regulation in the Induction of Trained Immunity. Semin Immunopathol, 2024. 46(3-4): p. 7.

37. Netea, M.G., et al., Defining trained immunity and its role in health and disease. Nat Rev Immunol, 2020. 20(6): p. 375–388.

38. Kim, H.Y. and W.W. Lee, Trained immunity induced by DAMPs and LAMPs in chronic inflammatory diseases. Exp Mol Med, 2025. 57(10): p. 2137–2147.

39. Jentho, E., et al., Trained innate immunity, long-lasting epigenetic modulation, and skewed myelopoiesis by heme. Proc Natl Acad Sci U S A, 2021. 118(42).

40. Sherwood, E.R., et al., Innate Immune Memory and the Host Response to Infection. J Immunol, 2022. 208(4): p. 785–792.

41. Amarasinghe, H.E., et al., Mapping the epigenomic landscape of human monocytes following innate immune activation reveals context-specific mechanisms driving endotoxin tolerance. BMC Genomics, 2023. 24(1): p. 595.

42. Li, X., et al., Histones: The critical players in innate immunity. Front Immunol, 2022. 13: p. 1030610.

43. Turner, B.M., Histone acetylation and control of gene expression. J Cell Sci, 1991. 99 (Pt 1): p. 13–20.

44. Tan, S.Y.X., J. Zhang, and W.W. Tee, Epigenetic Regulation of Inflammatory Signaling and Inflammation-Induced Cancer. Front Cell Dev Biol, 2022. 10: p. 931493.

45. Creyghton, M.P., et al., Histone H3K27ac separates active from poised enhancers and predicts developmental state. Proc Natl Acad Sci U S A, 2010. 107(50): p. 21931–6.

46. Song, H., et al., Histone post-translational modification and the DNA damage response. Genes Dis, 2023. 10(4): p. 1429–1444.

47. Ren, Y., et al., Histone deacetylase 8 regulates NF-kappaB-related inflammation in asthmatic mice through H3K9 acetylation. Chin Med J (Engl), 2022. 135(17): p. 2110–2112.

48. Bacher, S., et al., Regulation of Transcription Factor NF-kappaB in Its Natural Habitat: The Nucleus. Cells, 2021. 10(4).

49. Gruber, J.J., et al., HAT1 Coordinates Histone Production and Acetylation via H4 Promoter Binding. Mol Cell, 2019. 75(4): p. 711–724 e5.

50. Ortega, M.A., et al., Understanding HAT1: A Comprehensive Review of Noncanonical Roles and Connection with Disease. Genes (Basel), 2023. 14(4).

51. Liu, D., et al., Sepsis-induced immunosuppression: mechanisms, diagnosis and current treatment options. Mil Med Res, 2022. 9(1): p. 56.

52. Hotchkiss, R.S., G. Monneret, and D. Payen, Sepsis-induced immunosuppression: from cellular dysfunctions to immunotherapy. Nat Rev Immunol, 2013. 13(12): p. 862–74.

53. Mouton, W., et al., Towards standardization of immune functional assays. Clin Immunol, 2020. 210: p. 108312.

54. Vasuthas, K., et al., Type 1 diabetes mellitus (T1DM) does not affect whole blood responses to alginate-based microspheres despite plasma lipid and glucose differences. Mater Today Bio, 2025. 34: p. 102113.

55. Harm, S., et al., Blood Compatibility-An Important but Often Forgotten Aspect of the Characterization of Antimicrobial Peptides for Clinical Application. Int J Mol Sci, 2019. 20(21).

56. Yao, R.Q., et al., Advances in Immune Monitoring Approaches for Sepsis-Induced Immunosuppression. Front Immunol, 2022. 13: p. 891024.

57. Sehgal, R., et al., Granulocyte-Macrophage Colony-Stimulating Factor Modulates Myeloid-Derived Suppressor Cells and Treg Activity in Decompensated Cirrhotic Patients With Sepsis. Front Immunol, 2022. 13: p. 828949.

58. Chung, H., et al., Circulating Monocyte Counts and its Impact on Outcomes in Patients With Severe Sepsis Including Septic Shock. Shock, 2019. 51(4): p. 423–429.

59. Gao, Q., et al., Peripheral blood monocyte status is a predictor for judging occurrence and development on sepsis in older adult population: a case control study. BMC Emerg Med, 2023. 23(1): p. 11.

60. Li, Y., et al., Changes in Early T-Cell Subsets and Their Impact on Prognosis in Patients with Sepsis: A Single-Center Retrospective Study. Int J Clin Pract, 2023. 2023: p. 1688385.

61. Venet, F., et al., Immune Profiling Demonstrates a Common Immune Signature of Delayed Acquired Immunodeficiency in Patients With Various Etiologies of Severe Injury. Crit Care Med, 2022. 50(4): p. 565–575.

62. Ha, S.D., et al., HDAC8-mediated epigenetic reprogramming plays a key role in resistance to anthrax lethal toxin-induced pyroptosis in macrophages. J Immunol, 2014. 193(3): p. 1333–43.

63. Vaidya, K., et al., Identification of sequence determinants for the ABHD14 enzymes. Proteins, 2025. 93(1): p. 255–266.

64. Jo, C., et al., Histone acylation marks respond to metabolic perturbations and enable cellular adaptation. Exp Mol Med, 2020. 52(12): p. 2005–2019.

65. van der Knaap, J.A. and C.P. Verrijzer, Undercover: gene control by metabolites and metabolic enzymes. Genes Dev, 2016. 30(21): p. 2345–2369.

66. McBride, M.A., et al., The Metabolic Basis of Immune Dysfunction Following Sepsis and Trauma. Front Immunol, 2020. 11: p. 1043.

67. Liu, W., et al., Metabolic Reprogramming and Its Regulatory Mechanism in Sepsis-Mediated Inflammation. J Inflamm Res, 2023. 16: p. 1195–1207.

68. Foster, S.L., D.C. Hargreaves, and R. Medzhitov, Gene-specific control of inflammation by TLR-induced chromatin modifications. Nature, 2007. 447(7147): p. 972–8.

69. Novakovic, B., et al., *beta-Glucan Reverses the Epigenetic State of LPS-Induced Immunological Tolerance*. Cell, 2016. 167(5): p. 1354–1368 e14.

70. Saeed, S., et al., Epigenetic programming of monocyte-to-macrophage differentiation and trained innate immunity. Science, 2014. 345(6204): p. 1251086.

