## Supplemental data for "SLAMF1-peptide mediated epigenetic priming reprograms innate immune responses in sepsis"

### Supplementary materials

#### 1. Supplementary methods

**Gel-Based activity-based protein profiling assay to test Pen and P7-Pen compounds**

For the gel-based activity-based protein profiling assay (ABPP), ABHD14B WT and inactive S111A mutant [1, 2] recombinant proteins were treated with the peptides (Pen control peptide and P7-Pen), at 37 °C for 30 min with constant shaking (750 rpm) in a final volume of 100 μL. The protein concentration was constant (5 μM) and peptide concentration varied from 4 to 20 μM. After peptide treatment, 1 μM fluorophosphonate-rhodamine (FP-Rhodamine) activity probe (synthesized in-house) was added to the reaction mixture and incubated at 37 °C for 5 min with constant shaking, quenched by adding 4× SDS loading dye, and heated at 95 °C for 10 min. Prepared samples were loaded and resolved on a 12.5% SDS–PAGE gel, and the potential inhibitor activity of peptide was examined via in-gel fluorescence scanning using a iBright1500 gel documentation system (Invitrogen). To ensure the accurate protein loading in gels for ABPP assays, after scanning, the gels were stained with Coomassie Brilliant Blue R-250 and imaged using Syngene Chemi-XRQ gel documentation system.

**References**


1. Kumar, K., et al., *A Superfamily-wide Activity Atlas of Serine Hydrolases in Drosophila melanogaster.* Biochemistry, 2021. **60**(16): p. 1312–1324.
2. Rajendran, A., et al., *Functional Annotation of ABHD14B, an Orphan Serine Hydrolase Enzyme.* Biochemistry, 2020. **59**(2): p. 183–196.

#### 2. Supplementary Figures

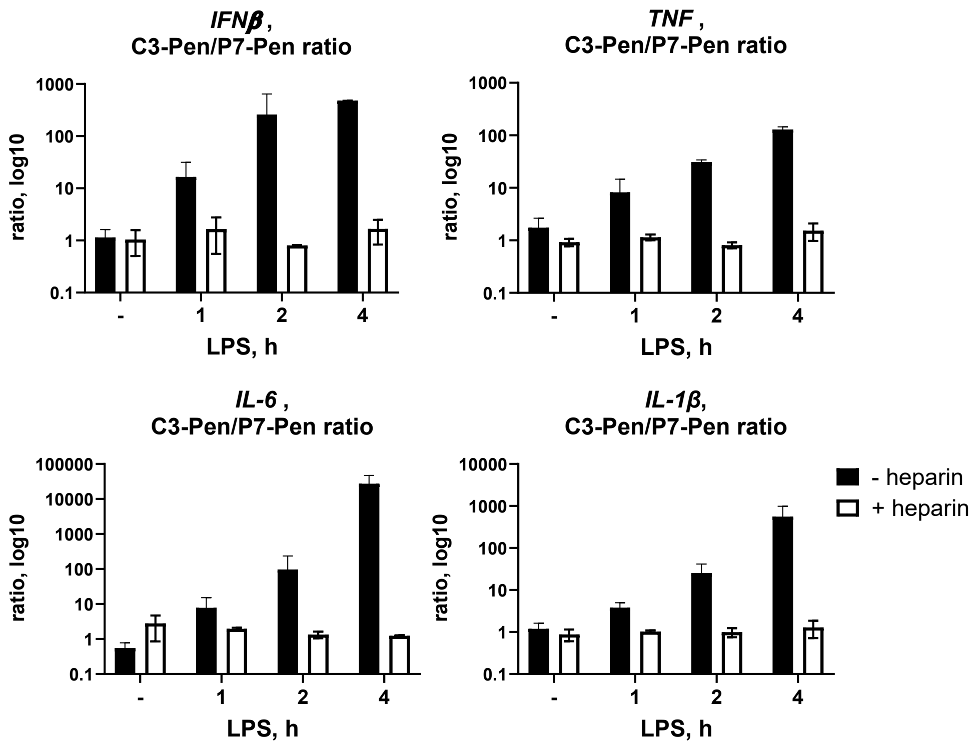


**Supplementary Figure 1. Heparin abrogates the inhibitory effect of P7-Pen on TLR4-induced cytokine expression.** Differentiated THP-1 cells were pretreated for 30 min with culture medium containing 15 μM control peptide (C3-Pen) or 15 μM P7-Pen, in the presence or absence of heparin (20 USP units/mL). Cells were subsequently stimulated with LPS (100 ng/mL) for the indicated time points, followed by RNA isolation and RT-qPCR analysis of cytokine gene expression. Graphs show RT-qPCR results from three independent experiments for *IFNB1*, *TNF*, *IL6*, and *IL1B* mRNA expression. Data were normalized to untreated controls and are presented as the ratio of P7-Pen to control peptide values on a logarithmic scale, expressed as mean relative fold change ± SD.

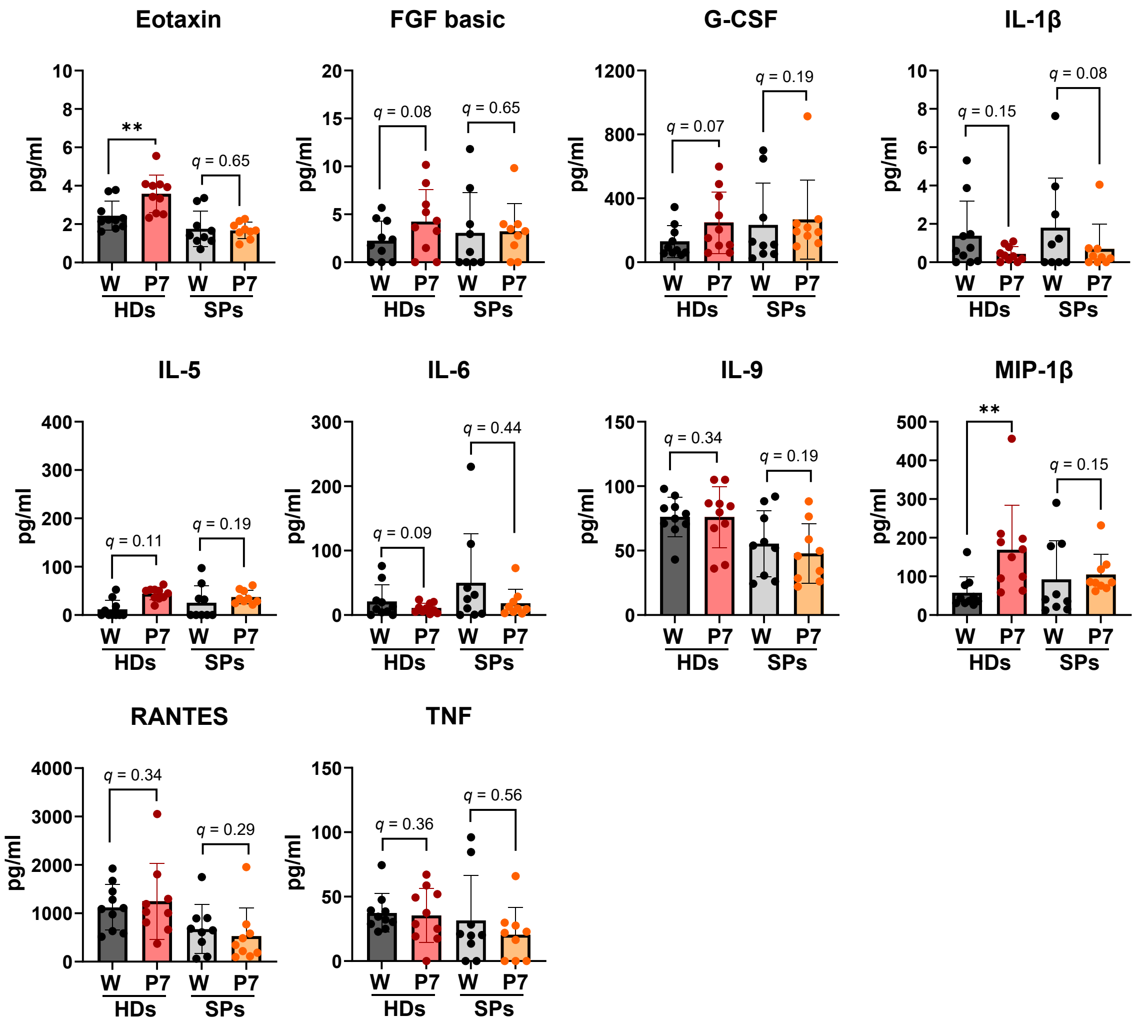


**Supplementary Figure 2. P7-Pen selectively modulates cytokine production in PBMCs from healthy donors and sepsis patients.** Graphs show cytokine secretion levels that were minimally affected or unchanged following P7-Pen treatment in PBMCs from HDs and SPs. Statistical significance was assessed using multiple unpaired *t*-tests across individual cytokines ($p < 0.05$, $*p < 0.01$, $**p < 0.001$).

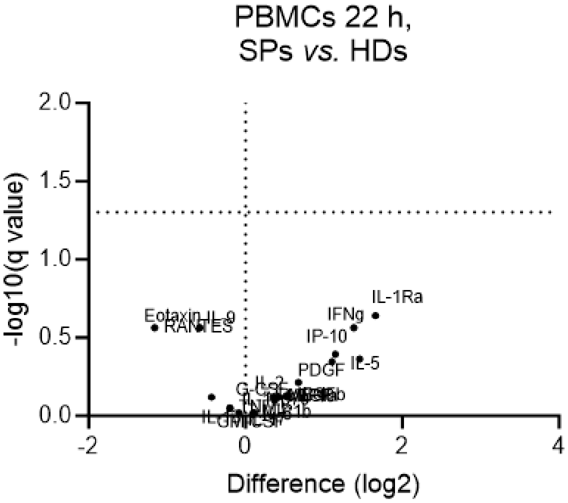


**Supplementary Figure 3. PBMCs from healthy donors and sepsis patients show no differences in cytokine secretion under control conditions.** PBMCs from healthy donors (HDs) and sepsis patients (SPs) were incubated for 20–22 h in medium containing vehicle (water). Volcano plot depicting cytokine secretion by PBMCs from SPs versus HDs, generated by plotting $\log_2$ fold change against $-\log_{10}$ FDR-adjusted *p* values ($q < 0.05$). Statistical significance for panels was assessed using multiple unpaired *t*-tests across individual cytokines.

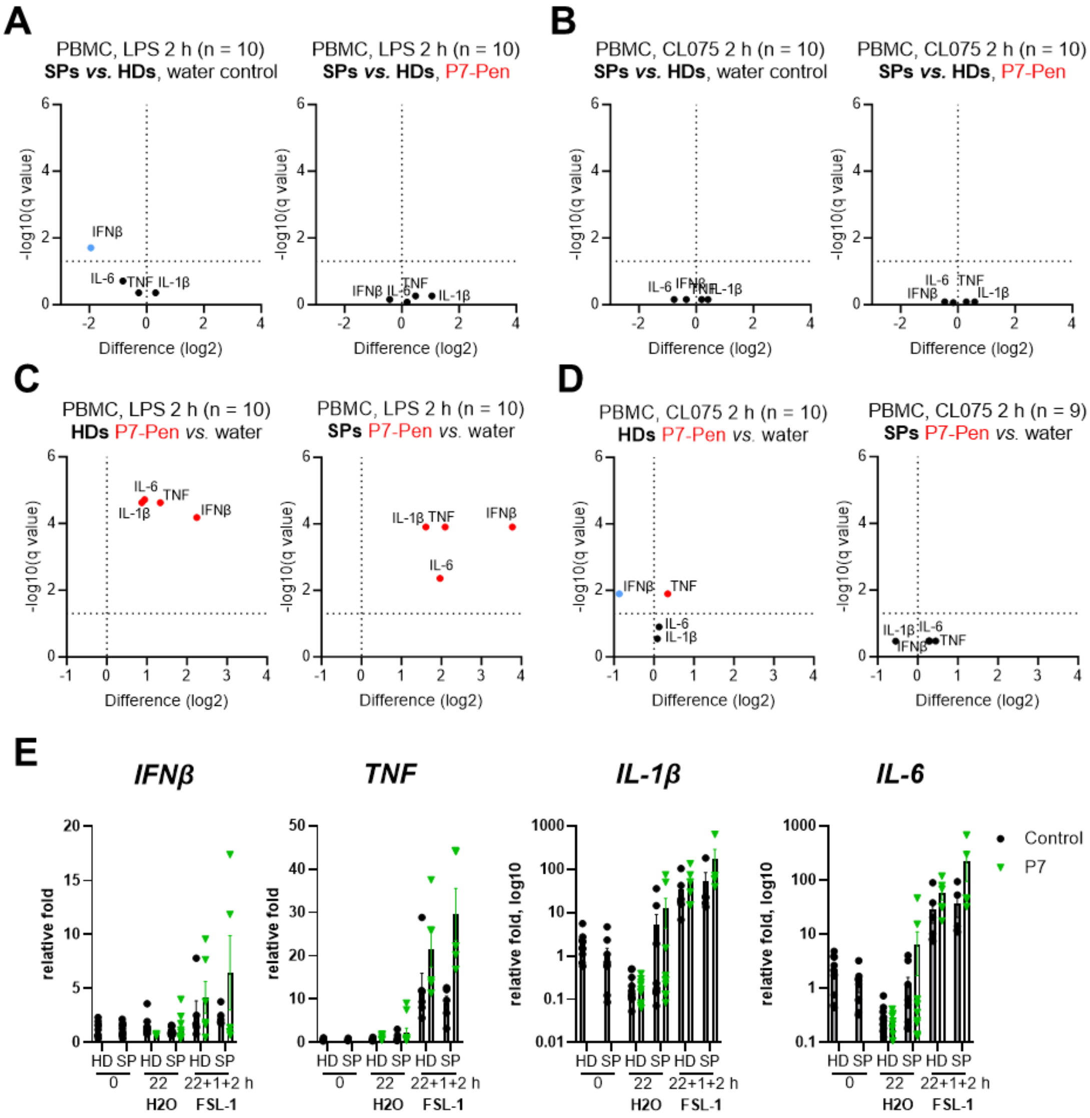


**Supplementary Figure 4. Priming of cells by P7-Pen enhances LPS-mediated cytokine mRNA expression in PBMCs from healthy donors and sepsis patients, with minimal effects on CL075-induced responses. (A, B)** Volcano plots showing comparison of relative cytokine mRNA induction between identical treatments in PBMCs from sepsis patients (SPs) and healthy donors (HDs) following stimulation with LPS (A) or CL075 (B) for 2 h. **(C, D)** Volcano plots comparing relative cytokine mRNA induction between P7-Pen–primed and control (water-treated) PBMCs from SPs and HDs following stimulation with LPS (C) or CL075 (D) for 2 h. **(A–D)** Cytokine mRNA levels were normalized to untreated samples and are presented as mean relative fold change ± SEM. Plots were generated by plotting $\log_2$ fold change against $-\log_{10}$ FDR-adjusted *p* values ($q < 0.05$). Statistical significance was assessed using multiple unpaired *t*-tests ($p < 0.05$,

*$p < 0.01$, **$p < 0.001$). **(E)** Relative expression levels of cytokine mRNA in PBMCs immediately after isolation (0), after 22 h incubation with vehicle (water), or following stimulation with FSL-1 for 2 h after 22 h priming with water or P7-Pen and a 1 h resting period (22 h + 1 h + 2 h).

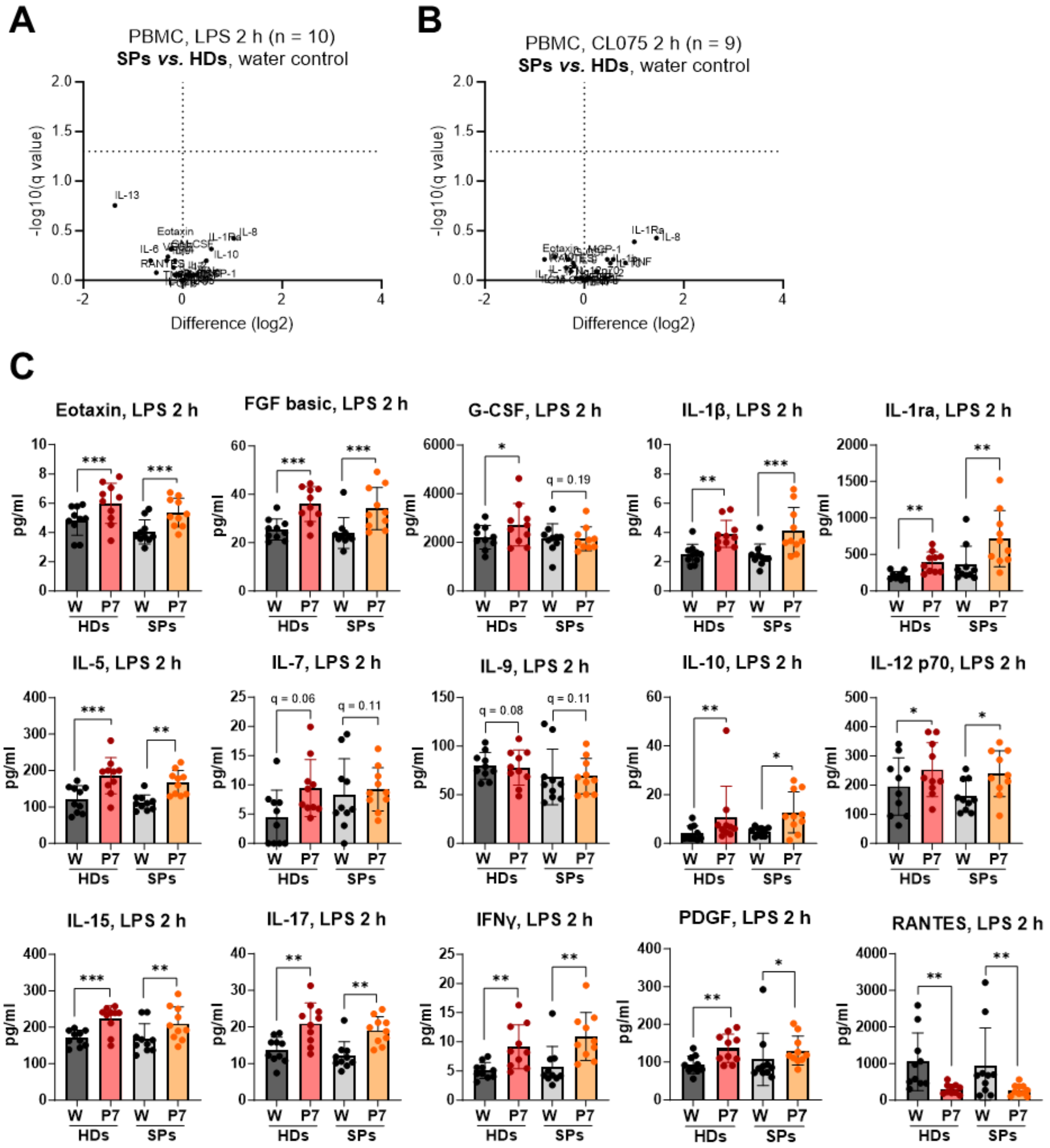


**Supplementary Figure 5. PBMCs from healthy donors and sepsis patients show no differences in cytokine secretion under control conditions, while P7-Pen priming enhances cytokine production following LPS stimulation. (A, B)** Volcano plots depicting cytokine secretion by PBMCs from sepsis patients (SPs) and healthy donors (HDs) primed with control medium (water) and stimulated with LPS (A) or CL075 (B). Plots were generated by plotting $\log_2$ fold change against $-\log_{10}$ FDR-adjusted $p$ values ($q < 0.05$). **(C)** Quantification of LPS-induced cytokine secretion in PBMCs primed with water or P7-Pen is shown as relative cytokine levels compared between treatment groups. Statistical significance for panels **(A–C)** was assessed using multiple unpaired *t*-tests ($p < 0.05$, *$p < 0.01$, **$p < 0.001$).

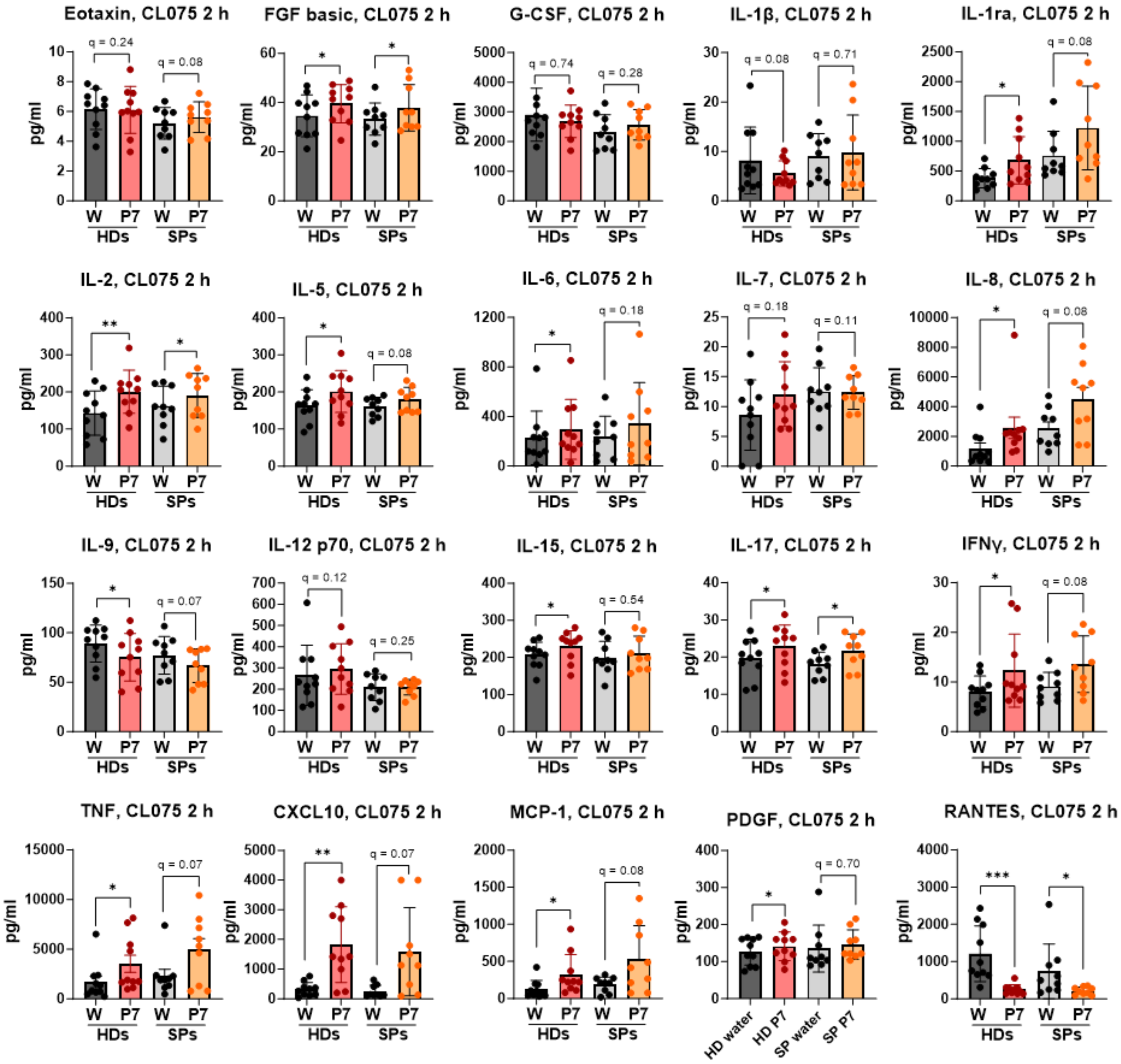


**Supplementary Figure 6. P7-Pen has minimal effects on cytokine secretion by PBMCs from healthy donors and sepsis patients following CL075 stimulation.** Graphs show cytokine secretion levels that were minimally affected or unchanged by P7-Pen treatment in PBMCs from healthy donors and sepsis patients. Statistical significance was assessed using multiple unpaired *t*-tests across individual cytokines ($p < 0.05$, *$p < 0.01$, **$p < 0.001$).

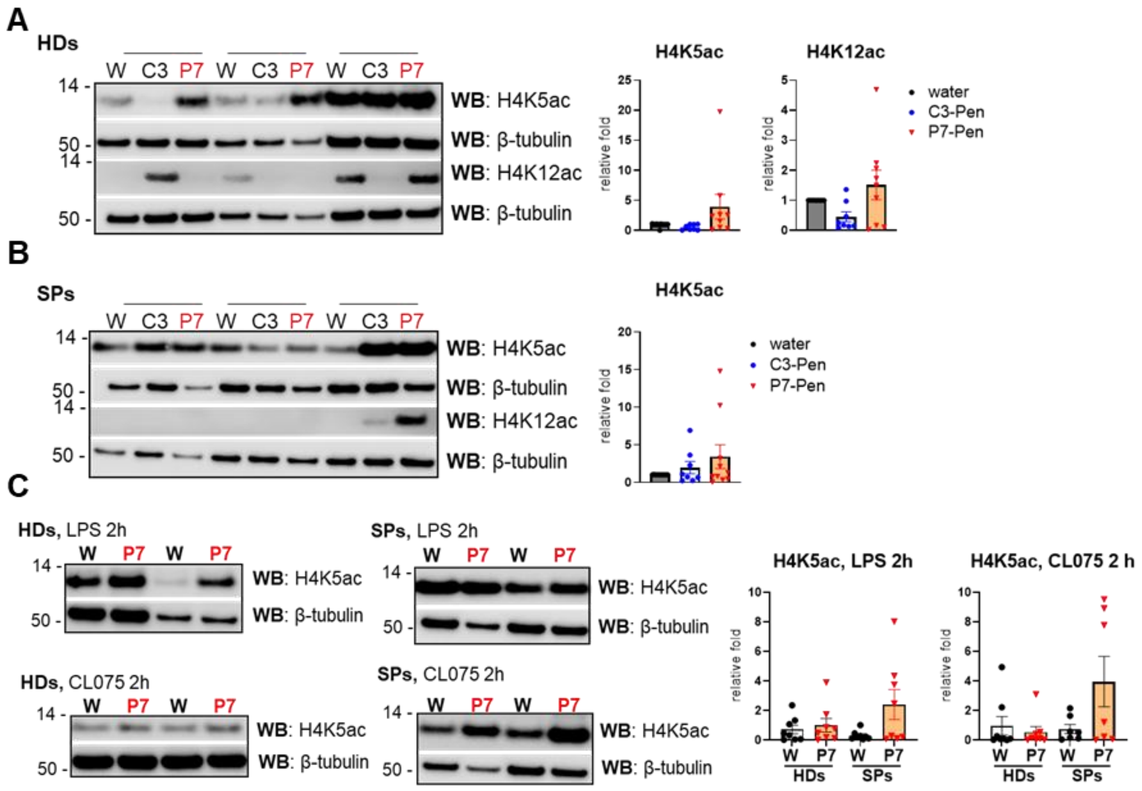


**Supplementary Figure 7. P7-Pen priming does not significantly alter histone H4 acetylation levels in PBMCs following TLR stimulation.** Western blot analysis illustrates histone H4 acetylation patterns together with quantitative densitometry for all analyzed samples. **(A, B)** Representative immunoblots of PBMC lysates from healthy donors (A) or sepsis patients (B) showing levels of H4K5ac and H4K12ac following priming with vehicle (water), 15 µM control peptide C3-Pen, or 15 µM P7-Pen for 20–22 h. **(C)** Representative immunoblots (2 of 10 donors per group) showing H4K5ac levels in PBMCs primed with water or P7-Pen and subsequently stimulated with LPS or CL075 for 2 h. Quantification shown in the graphs represents signal intensity normalized to the corresponding loading control (β-tubulin). Data are presented as mean relative fold change ± SEM. Statistical significance was assessed using the nonparametric Wilcoxon matched-pairs signed-rank test ($p < 0.05$).

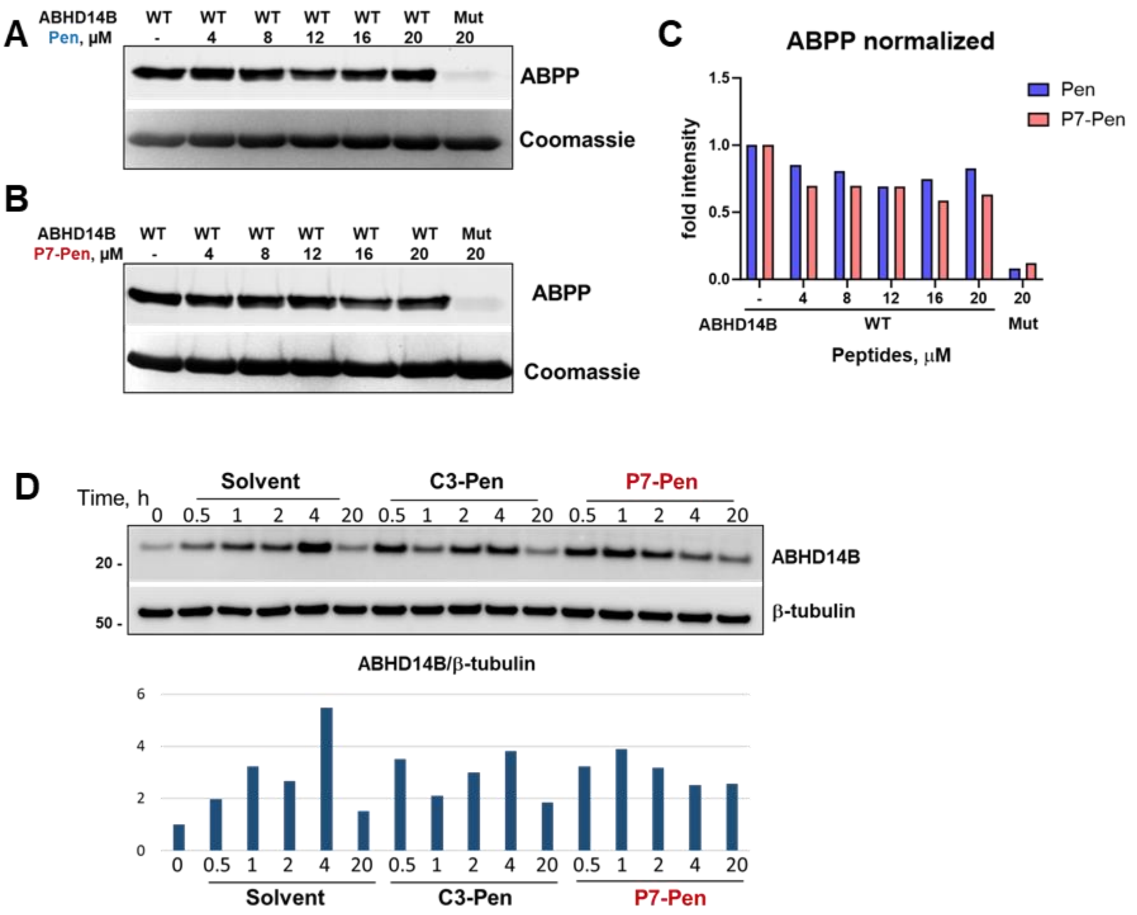


**Supplementary Figure 8. Activity-based protein profiling demonstrates that P7-Pen does not inhibit the catalytic activity of ABHD14B.** (**A–C**) Purified recombinant ABHD14B wild-type (WT) protein and the catalytically inactive ABHD14B S111A mutant were pre-incubated with increasing concentrations of control peptide (Pen), P7-Pen, or vehicle control at 37 °C for 30 min, followed by labeling with the activity-based probe FP-Rhodamine for 5 min. Fluorescence imaging shows FP-Rhodamine–labeled ABHD14B in samples treated with Pen or P7-Pen, while the S111A mutant served as a negative control. Corresponding Coomassie Brilliant Blue–stained gels confirm equal protein loading, and quantification of FP-Rhodamine signal normalized to total protein is shown as fold change relative to control. (**D**) The effect of P7-Pen on ABHD14B total protein level was further assessed by Western blot analysis of PBMCs treated with vehicle (water), 15 μM control peptide (C3-Pen), or 15 μM P7-Pen for indicated time. Representative blots are from one of three independent donors. Densitometric analyses for ABHD14B signal intensity normalized to the loading control β-tubulin in representative image is shown on graph under the blots.
